# Metabolic Adaptations Rewire CD4 T Cells in a Subset-Specific Manner in Human Critical Illness with and without Sepsis

**DOI:** 10.1101/2025.01.27.635146

**Authors:** Matthew T. Stier, Allison E. Sewell, Erin L. Mwizerwa, Chooi Ying Sim, Samantha M. Tanner, Casey M. Nichols, Heather H. Durai, Erin Q. Jennings, Paul Lindau, Erin M. Wilfong, Dawn C. Newcomb, Julie A. Bastarache, Lorraine B. Ware, Jeffrey C. Rathmell

**Affiliations:** Division of Allergy, Pulmonary & Critical Care Medicine, Department of Medicine, Vanderbilt University Medical Center, Nashville, TN, United States; Vanderbilt Center for Immunobiology, Vanderbilt University Medical Center, Nashville, TN, United States; Department of Medicine, Vanderbilt University Medical Center, Nashville, TN, United States.; Department of Pathology, Microbiology, and Immunology, Vanderbilt University Medical Center, Nashville, TN, United States; Division of Hematology and Oncology, Department of Medicine, Vanderbilt University Medical Center, Nashville, TN, United States; Division of Rheumatology and Immunology, Department of Medicine, Vanderbilt University Medical Center, Nashville, TN, United States; Department of Cell and Developmental Biology, Vanderbilt University, Nashville, TN, United States

**Author notes:** Corresponding Author: Jeffrey C. Rathmell, PhD.

## Abstract

Host immunity in sepsis has features of hyperinflammation together with progressive immunosuppression, particularly among CD4 T cells, that can predispose to secondary infections and ineffectual organ recovery. Metabolic and immunologic dysfunction are archetypal findings in critically ill patients with sepsis, but whether these factors are mechanistically linked remains incompletely defined. We characterized functional metabolic properties of human CD4 T cells from critically ill patients with and without sepsis and healthy adults. CD4 T cells in critical illness showed increased subset-specific metabolic plasticity, with regulatory T cells (Tregs) acquiring glycolytic capacity that stabilized suppressive markers FOXP3 and TIGIT and correlated with clinical illness severity. Single-cell transcriptomics identified differential kynurenine metabolism in Tregs, which was validated *ex vivo* as a mechanism of Treg glycolytic adaptation and suppressive rewiring. These findings underscore immunometabolic dysfunction as a driver of CD4 T cell remodeling in sepsis and suggest therapeutic avenues to restore an effective immune response.

## INTRODUCTION

Critical illness requiring intensive care unit (ICU) admission and life support due to severe single or multiorgan failure affects over 5 million patients annually in the United States^1^. Among the most common ICU diagnoses is sepsis, a dysregulated host response to infection that leads to end-organ injury^2^. Sepsis accounts domestically for over 200,000 deaths per year and more than $24 billion in healthcare expenditures, making it the costliest condition treated in U.S. hospitals^3,4^. Current management relies on antimicrobials and/or surgical source control to address the inciting infection alongside supportive care measures—including vasopressors, mechanical ventilation, renal replacement therapy, and blood product transfusions—to sustain organ function and allow for natural repair and recovery. Despite decades of research, no targeted therapies have been developed to effectively treat the pathophysiology of sepsis^5–7^.

Immunologic dysfunction is a hallmark of sepsis, encompassing both hyperinflammatory and immunosuppressive states. Early sepsis is characterized by excessive inflammatory responses, including elevated levels of pro-inflammatory cytokines such as TNF-α and IL-1β (e.g., “cytokine storm”), which contribute to tissue injury and organ failure^8^. In parallel, a compensatory but disproportionate immunosuppressive response, often referred to as “immunoparalysis”, develops and can persist past the acute phase. This immunosuppressive state predisposes patients to secondary infections, persistent organ dysfunction, hospital readmissions, and death^8–19^. CD4 T cells are central to this immunosuppressive remodeling, evidenced by an “exhausted-like” phenotype with decreased effector function^9,20–31^. The severity of this cell-mediated immune impairment is underscored by the frequent reactivation of latent viruses (e.g. CMV, EBV, HSV) in critically ill patients^32–34^, occurring at rates comparable to those observed in solid organ transplant recipients receiving T cell immunosuppressants^35,36^. While preclinical animal studies have sought to uncover mechanisms influencing CD4 effector and regulatory T cell (Treg) function in sepsis, these findings have yet to yield clinically actionable outcomes or robust human validation, leaving a mechanistic gap in our understanding of CD4 T cell remodeling in critically ill and septic patients.

Metabolic dysfunction is another well-recognized feature of critical illness including sepsis^37,38^. Immunometabolism, defined as the metabolic processes of immune cells, is a mechanistic determinant of their cell states and functions, and can play both a protective or pathologic role in conditions including obesity, autoimmunity, cancer, and SARS-CoV-2 infection^39,40^. Accordingly, metabolic reprogramming has emerged as a promising therapeutic target with numerous approaches currently in clinical trials, particularly in autoimmunity and cancer^41,42^. While prior studies in human sepsis have reported abnormalities in mitochondrial complex I/V activities and broad impairments in glycolysis and oxidative phosphorylation among bulk T lymphocytes^43,44^, the overall role of CD4 T cell immunometabolism remains understudied. It is also unclear whether sepsis uniquely drives these metabolic changes or whether they are broadly reflective of critical illness. In this study, we used biospecimens from critically ill patients to comprehensively profile the metabolic dependencies and capacities of major CD4 T cell subsets. We aimed to determine whether metabolic alterations drive CD4 T cell remodeling toward immunosuppression and associate with poor clinical outcomes, thereby providing key insights into immunometabolic dysfunction during critical illness with and without sepsis. Our findings revealed that heightened metabolic plasticity via differential glycolytic and kynurenine metabolism promoted Treg resilience and suppressive function at the expense of conventional CD4 T cell subsets.

## RESULTS

### Metabolic Phenotyping of CD4 T Cells in Critical Illness

We prospectively collected peripheral blood mononuclear cells (PBMC) from consented patients in the medical and surgical intensive care units (ICUs) at Vanderbilt University Medical Center (Fig. 1a and Extended Data Table 1). We adjudicated these into critically ill patients that were non-septic (CI-NS) and critically ill patients with sepsis (CI-Sep) (Extended Data Fig. 1). Complete blood counts from these participants showed a mixed neutrophilic/monocytic response with marked lymphopenia, most notable in CI-Sep in comparison to community-matched non-acutely ill healthy controls (NHC), consistent with prior reports (Fig. 1b,c).

**Figure 1:**
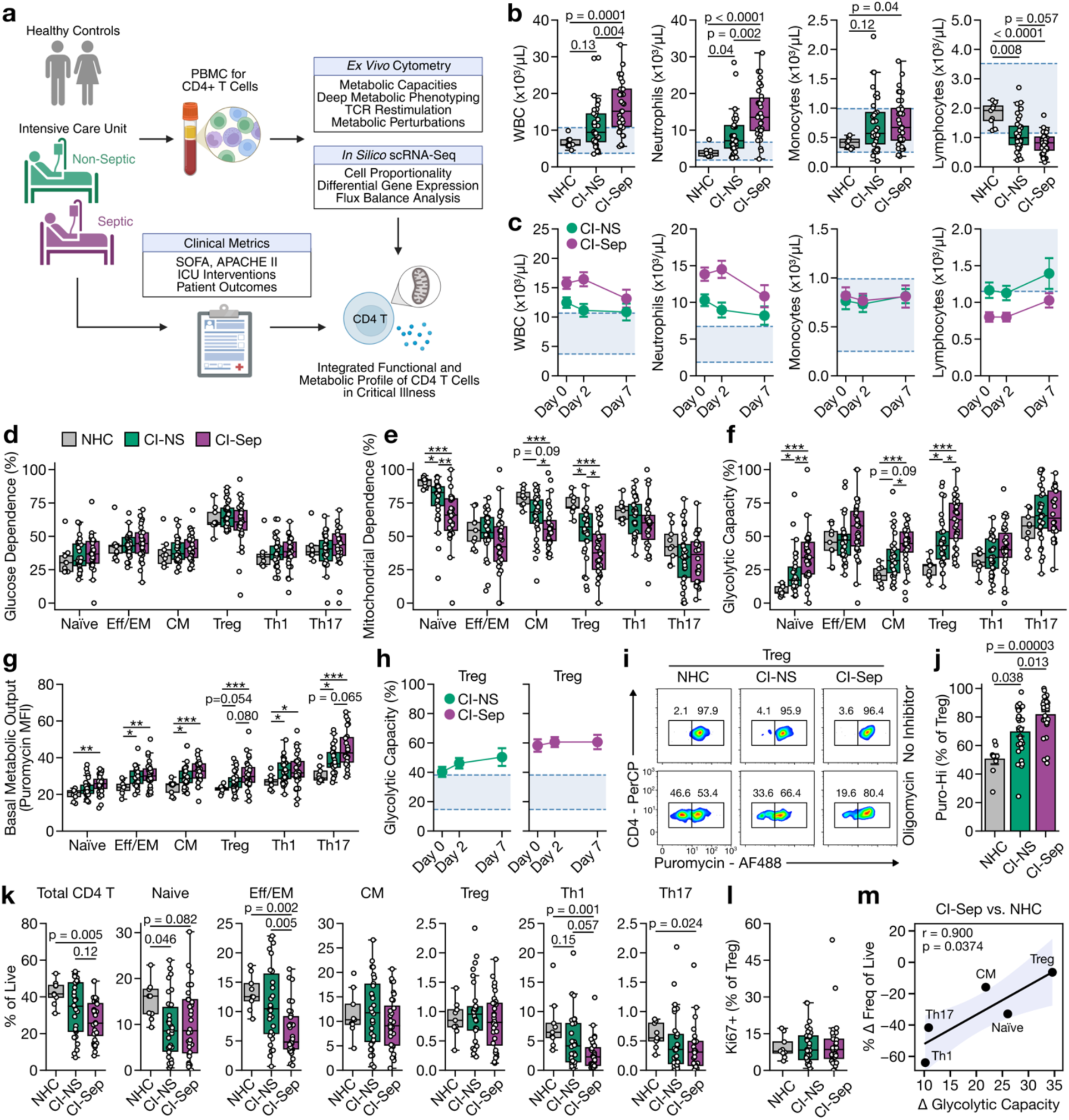
CD4 T cells exhibit subset-specific metabolic adaptations in the context of critical illness and severe sepsis. **a,** Project overview. Panel created in BioRender.com. **b,** Differential cell counts at day 2 post-ICU admission. **c**, Differential cell counts from ICU admission (day 0) and subsequently on day 2 and 7 post-ICU admission. Shaded region represents the assay normal reference range. **d,e,f,** Glucose dependence, mitochondrial dependence, and glycolytic capacity per CD4 T cell subset determined by puromycin-incorporation assay at day 2 post-ICU admission. Displayed as percentage of the maximum puromycin incorporation without metabolic inhibitors. **g,** Basal metabolic output as determined by the geometric MFI of puromycin in the absence of metabolic inhibitors. **h,** Time course of glycolytic capacity in Treg at day 0, 2, and 7 post-ICU admission. Shaded region represents an estimated normal range using NHC Treg from **c** and plotting the mean ± 2 standard deviations. **i,j** Representative flow plots and summarized data for puromycin incorporation in the setting of either no inhibitor or with oligomycin at day 2 post-ICU admission. Boxes denoted Puro-Hi and Puro-Lo populations. Percentages are of the Treg parent gate. **k,** Frequency of CD4 T cell subsets as a percentage of all live cells at day 2 post-ICU admission. **l,** Percentage of Ki67+ Treg among all Treg at day 2 post-ICU admission. **m,** Spearman rank correlation with shaded confidence interval among CD4 T cell subsets between the percentage change in the population mean frequency of live cells in CI-Sep compared to NHC versus the population mean change in glycolytic capacity. Statistical analysis for **b, d-g, j-I** was performed using the Kruskal-Wallis test with Dunn’s post-hoc testing. Where able, actual p values are shown. In certain instances, due to the density of the data and labels, p values are reflected as: *p < 0.05, **p < 0.01, and ***p < 0.001. Error bars for **b,d-g,k,l** represent 95% confidence intervals and for **c,h,j** represent SEM. WBC = White blood cell. K-W = Kruskal-Wallis.

To decipher the immunometabolic landscape of CD4 T cells in critical illness with and without sepsis, we performed functional assessments of metabolic dependencies and capacities using a puromycin incorporation assay^45^. We focused on samples collected between 48-72 hours (i.e. day 2) after ICU admission, where both inflammatory and immunosuppressive mechanisms are expected to be active. CD4 T cell subsets including naïve, effector/effector memory (Eff/EM), central memory (CM), regulatory T cells (Treg), and effector subsets T helper 1 and 17 cells (Th1, Th17) exhibited no difference in their metabolic dependence on glucose, fatty acid oxidation, and glutaminolysis (Fig. 1d and Extended Data Fig. 2). However, there were significant subset-specific differences in mitochondrial dependence and acquired glycolytic capacity between the critically ill cohort and NHC (Fig. 1e,f). Specifically, Treg show the most substantial acquisition of glycolytic capacity while the effector lineages (Eff/EM, Th1, Th17) demonstrated the least. This occurred in the context of broadly increased metabolic output across all CD4 T cell subsets in critical illness compared to NHC (Fig. 1g). These data suggested that while Treg do not exhibit a higher baseline dependence on glucose (Fig. 1d), their increased glycolytic capacity (Fig. 1f) reflects a metabolic plasticity that may enable sustained ATP production and suppressive functions under the mitochondrial stress conditions present in critically ill patients. Notably, these observed differences were magnified in CI-Sep compared to CI-NS. Additionally, in a subset of participants where serially collected samples were available, this Treg acquired glycolytic capacity increased (CI-NS) or remained persistently elevated (CI-Sep) over the first week of their hospitalization (Fig. 1h). Furthermore, while two metabolically distinct programs were apparent among Treg in NHC marked by sensitivity (Puro-Lo) or resistance (Puro-Hi) to the mitochondrial ATPase inhibitor oligomycin (Fig. 1i) the oligomycin-resistant Puro-Hi population was considerably enriched in CI-Sep (Fig. 1i,j). This oligomycin resistance is consistent with elevated glycolytic capacity and metabolic plasticity.

While we observed lymphopenia broadly in CI-NS and CI-Sep, the degree of attrition was variable across CD4 T cell subsets (Fig. 1k). Naïve and effector lineages (Eff/EM, Th1, and Th17) had reduced frequency particularly in CI-Sep compared to NHC, whereas Treg and CM frequencies were preserved. These findings align with prior studies showing that stable or increased Treg frequencies in septic patients result from disproportionate loss of conventional CD4 T cell subsets^24,25,31^ and were not explained by increased Treg proliferation compared to NHC (Fig. 1l). Strikingly, there was a positive correlation among CD4 T cell subsets between the magnitude of acquired glycolytic capacity and their change in frequency in CI-Sep compared to NHC (Fig. 1m). These data suggest that metabolic plasticity in the form of acquired glycolytic capacity provides a cell survival advantage in critical illness, which is most substantial in the Treg lineage contributing to the progressive development of immunosuppression.

We further characterized the glycolytic and oxidative attributes of these CD4 T cell subsets. CI-Sep Treg expressed higher levels of the glycolysis rate-limiting enzyme hexokinase 1 (HK1) compared to NHC, though this was also observed in naïve, Eff/EM, and CM populations suggesting that this may be necessary but not sufficient for acquired glycolytic capacity in critical illness (Fig. 2a). However, not all glycolytic enzymes were upregulated as evidence by stable levels of GAPDH in CI-NS and CI-Sep (Fig. 2b). We next assessed mitochondrial mass and membrane potential (MMP), noting that naïve CD4 T cells have unique changes in their mitochondrial attributes with overall higher mass, MMP, and ratio of MMP:mass in CI-Sep compared to NHC (Fig. 2c-g). Interestingly, there was an increase in Eff/EM cells in CI-Sep with high mitochondrial mass but negligible MMP (Mass-Hi MMP-Lo), which is frequently observed in cells with dysfunctional mitochondria or undergoing programmed cell death (Fig. 2g,h). This is consistent with the high rate of attrition of Eff/EM cells in critical illness with and without sepsis (Fig. 1k). Finally, we observed a tendency for higher levels of mitochondrial stress in the form of mitochondrial reactive oxygen species by MitoSOX staining across all CD4 T cell subsets in CI-Sep compared to NHC (Fig. 2j).

**Figure 2:**
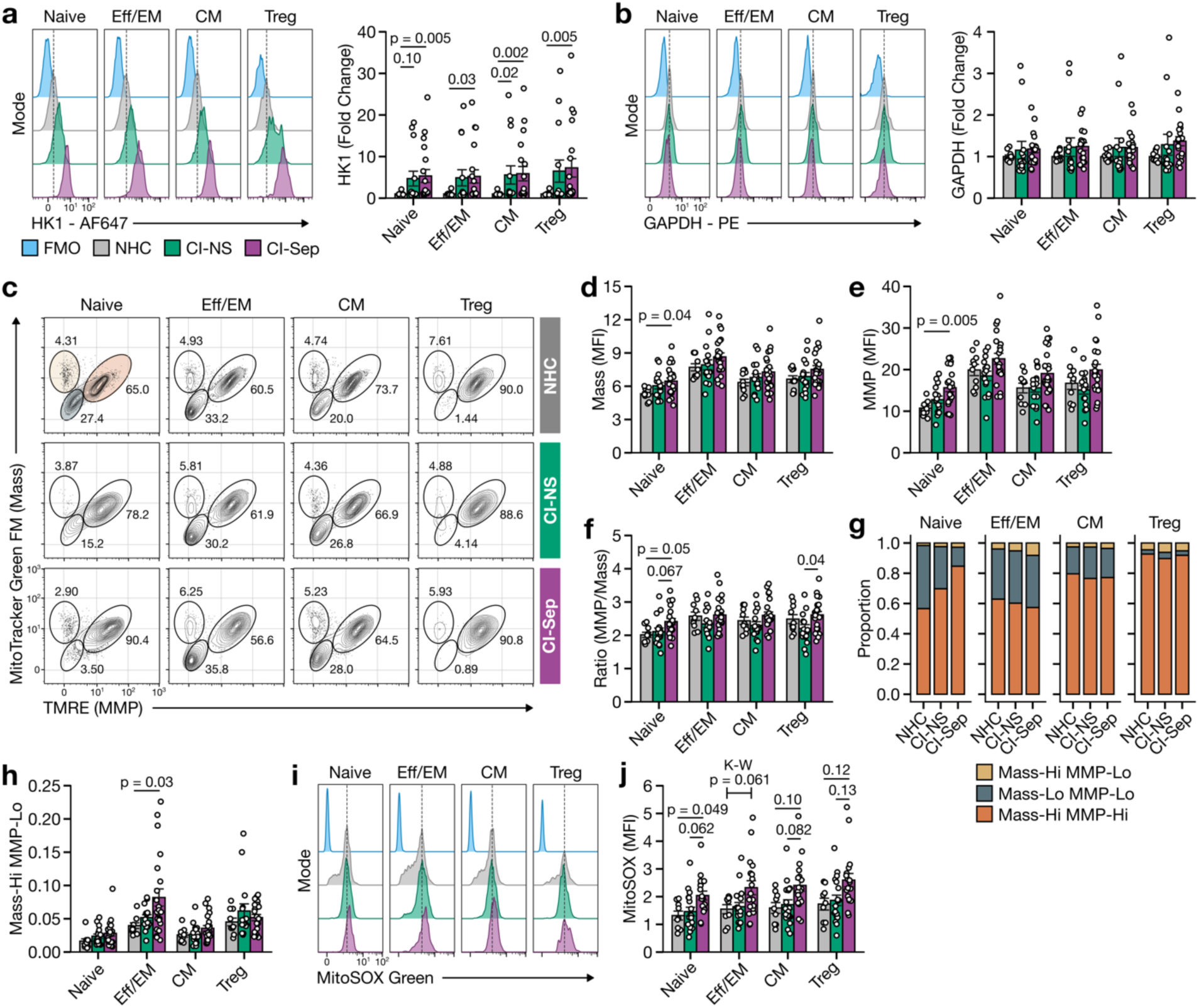
Glycolytic and mitochondrial characteristics of CD4 T cell subsets in critical illness and severe sepsis. **a,b,** Protein expression of hexokinase 1 (HK1) and glyceraldehyde 3-phosphate dehydrogenase (GAPDH) by geometric MFI normalized to fold change of NHC. **c,** Representative flow cytometry plots of mitochondrial mass measured by MitoTracker Green FM and mitochondrial membrane potential (MMP) measured by TMRE. **d,e,** Summary of mitochondrial mass (geometric MFI of MitoTracker Green FM) and MMP (geometric MFI on TMRE) among mass-hi/MMP-hi cells (orange-shaded gate). **f,** Ratio of MMP:mitochondrial mass. **g**, Proportion of cells in **c-f** represented by mitochondrial mass-hi/MMP-hi (orange-shaded gate), mitochondrial mass-lo/MMP-lo (steel blue-shaded gate), and mitochondrial mass-hi/MMP-lo (yellow-shaded gate). **h,** Bar and strip plot representation from **g** of the mitochondrial mass-hi/MMP-lo group. **i,j,** Mitochondrial reactive oxygen species as measured by the geometric MFI of MitoSOX Green. Statistical analysis for **a,b,d-f,h,j** was performed using the Kruskal-Wallis test followed by Dunn’s post-hoc testing. FMO = fluorescence minus one control; MFI = mean fluorescence intensity (geometric).

### Functional Characterization of CD4 T Cells in Critical Illness

We next assessed the functional capacities of conventional CD4 T cells (Tconv) including naïve, Eff/EM, and CM as well as CD4 Treg. PBMC were restimulated with T cell receptor-directed stimulation (anti-CD3/CD28/CD2) for 24 hours with subsequent flow cytometric assessment of cytokines and functional markers on CD4 Tconv and Treg. In Tconv from CI-Sep, we observed an increased frequency of TNFα-, IL-17A-, and IL-4-producing cells, but not IFNγ-producing cells, compared to NHC (Fig. 3a,b and Extended Data Fig. 3a). However, TNFα expression was reduced on a per-cell basis in CI-Sep, consistent with a more exhausted-like phenotype (Fig. 3b). IL-2-expressing Tconv were also more frequent in CI-NS and CI-Sep, potentially reflecting an autocrine and/or paracrine survival response (Fig. 3b). Tconv in CI-NS and CI-Sep exhibited increased frequency and expression of activation (CD25) and checkpoint inhibitor (PD-1 and LAG-3) proteins, suggesting a mixed phenotype of activation and exhaustion consistent with their cytokine profile (Fig. 3c).

**Figure 3:**
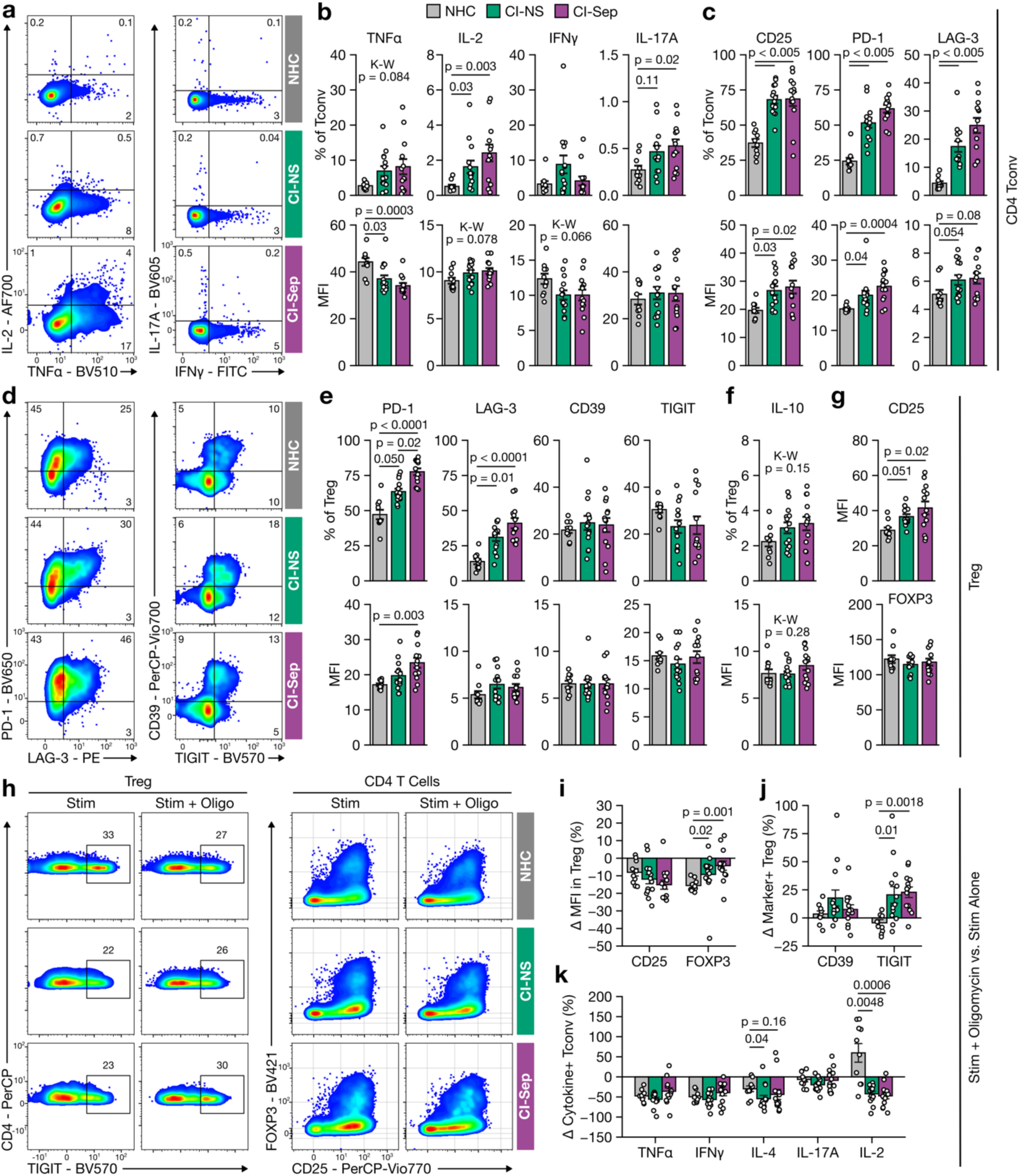
Mitochondrial stress triggers altered functionality in CD4 T cells. **a,b** Representative flow plots and summary data of conventional T-helper cytokine protein expression (TNFα, IL-2, IFNγ, and IL-17A) in Tconv (CD4+ FOXP3-) of PBMC restimulated for 24 hrs with anti-CD3/CD28/CD2. Both cytokine-positive percentage of Tconv (top panels) and geometric MFI of the listed cytokine among cytokine-positive cells (bottom panels) is shown. **c,** Protein expression of CD25, PD-1, and LAG-3 on Tconv with both percent-positive (top panels) and geometric MFI (bottom panels) shown. **d,e,f** Representative flow plots and summary data of suppressive marker and cytokine protein expression on Treg (CD4+ FOXP3+) with both percent-positive (top panels) and geometric MFI (bottom panels) shown. **g,** Protein expression of CD25 (top panel) and FOXP3 (bottom panel) on Treg by geometric MFI. **h,** Representative flow plots of TIGIT protein expression on Treg and FOXP3/CD25 protein expression marking Treg among CD4+ T cells in anti-CD3/CD28/CD2 restimulated PBMC with stimulation alone or with stimulation plus sublethal mitochondrial disruption with oligomycin (10 nM). **i** Summary data for CD25 and FOXP3 protein expression from **h** represented as a percentage change in geometric MFI among stimulated, oligomycin-treated Treg compared to stimulated-only. **j,** Summary data for CD39+ Treg and TIGIT+ Treg from **h** represented as a percentage change in the frequency of CD39+ Treg and TIGIT+ Treg from stimulated, oligomycin-treated compared to stimulated-only. **k,** Summary data for cytokine-positive Tconv from **h** represented as a percentage change in the frequency of cytokine-positive Tconv from stimulated, oligomycin-treated compared to stimulated-only. Statistical analysis for **b,c,e-g, i-k** was performed using the Kruskal-Wallis test followed by Dunn’s post-hoc testing. K-W = Kruskal-Wallis; MFI = mean fluorescence intensity (geometric).

Tregs also exhibited functional and phenotypic differences in CI-NS and CI-Sep compared to NHC, with a higher frequency and expression of PD-1 and LAG-3, markers that are associated with suppressive effector Tregs in humans (Fig. 3d,e)^46–48^. Tregs in NHC, CI-NS, and CI-Sep expressed comparable frequency and expression of CD39 and TIGIT (Fig. 3d,e). There was also no difference in the frequency of IL-10-producing Tregs, IL-10 expression levels, or in the acquisition of pro-inflammatory capacities (Fig. 3f and Extended Data Fig. 3b). Moreover, CI- Sep and CI-NS Tregs had increased CD25 expression, which is associated with suppressive Treg^49^, and stable levels of the canonical Treg transcription factor FOXP3 compared to NHC Treg (Fig. 3g). Overall, Treg in critically ill patients demonstrated a suppressive phenotype.

The enhanced glycolytic capacity in CD4 T cells may allow functional adaptation in the setting of mitochondrial stress, which is systemic in critical illness states but rarely modeled in functional analyses of human leukocytes from ICU patients. To investigate this key interplay, we next examined how mitochondrial disruption impacts the phenotypic and functional properties of Tconv and Treg subsets. We restimulated PBMCs as previously described with or without a sublethal dose of oligomycin (10 nM) to partially inhibit mitochondrial energetics. Mitochondrial stress diminished FOXP3 expression in NHC Treg whereas FOXP3 expression was relatively preserved in CI-NS and CI-Sep Treg (Fig. 3h,i). Additionally, the frequency of TIGIT-expressing Treg increased in CI-NS and CI-Sep Treg exposed to mitochondrial stress, which was not observed in NHC Treg (Fig. 3h,j and Extended Data Fig. 3c). Prior work has shown that FOXP3 expression levels correlate with Treg suppressor activity^50,51^ and TIGIT+ Treg are potent suppressors of Th1 and Th17 responses^52^. Other markers of Treg function including CD25, CD39, and IL-10 were similarly altered by mitochondrial stress among the groups (Fig. 3h-j and Extended Data Fig. 3c-d). Treg immune checkpoint inhibitor expression data was mixed in the presence of mitochondrial stress, with disproportionate reductions in LAG-3 but not PD-1 positivity in CI-NS and CI-Sep compared to NHC (Extended Data Fig. 3e). Collectively, our data highlight that CI-NS and CI-Sep Treg adopt a more stable suppressive phenotype compared to NHC Treg when challenged with mitochondrial stress.

Among Tconv, mitochondrial stress universally reduced the frequency of TNFα-, IFNγ-, IL-4-, and IL-17A-expressing cells across NHC, CI-NS, and CI-Sep (Fig. 3k and Extended Data Fig. 3f). Mitochondrial stress was associated with a larger decrease in Tconv PD-1 and LAG-3 expression in CI-NS and CI-Sep compared to NHC (Extended Data Fig. 3g). Strikingly, CI-NS and CI-Sep Tconv exhibited a reduction in IL-2 expression compared to NHC Tconv (Fig. 3k). This impaired production of IL-2, a key pro-survival cytokine in T cells, highlights a metabolic vulnerability in Tconv from critically ill patients, which may underlie their disproportionate attrition during critical illness with and without sepsis.

### Transcriptomic Map of Critical Illness

To investigate the mechanisms underlying metabolic remodeling in CD4 T cells, we performed single-cell RNA sequencing (scRNA-seq) on PBMCs from 9 NHC, 19 CI-NS, and 19 CI-Sep patients. After quality control and preprocessing, we generated a dataset comprised of 644,147 individual cells. We annotated 27 distinct PBMC lineages using canonical transcriptional markers, with sufficient resolution to identify even rare populations, including non-NK innate lymphoid cells (ILC) (Fig. 4a,b and Extended Data Fig. 4).

**Figure 4:**
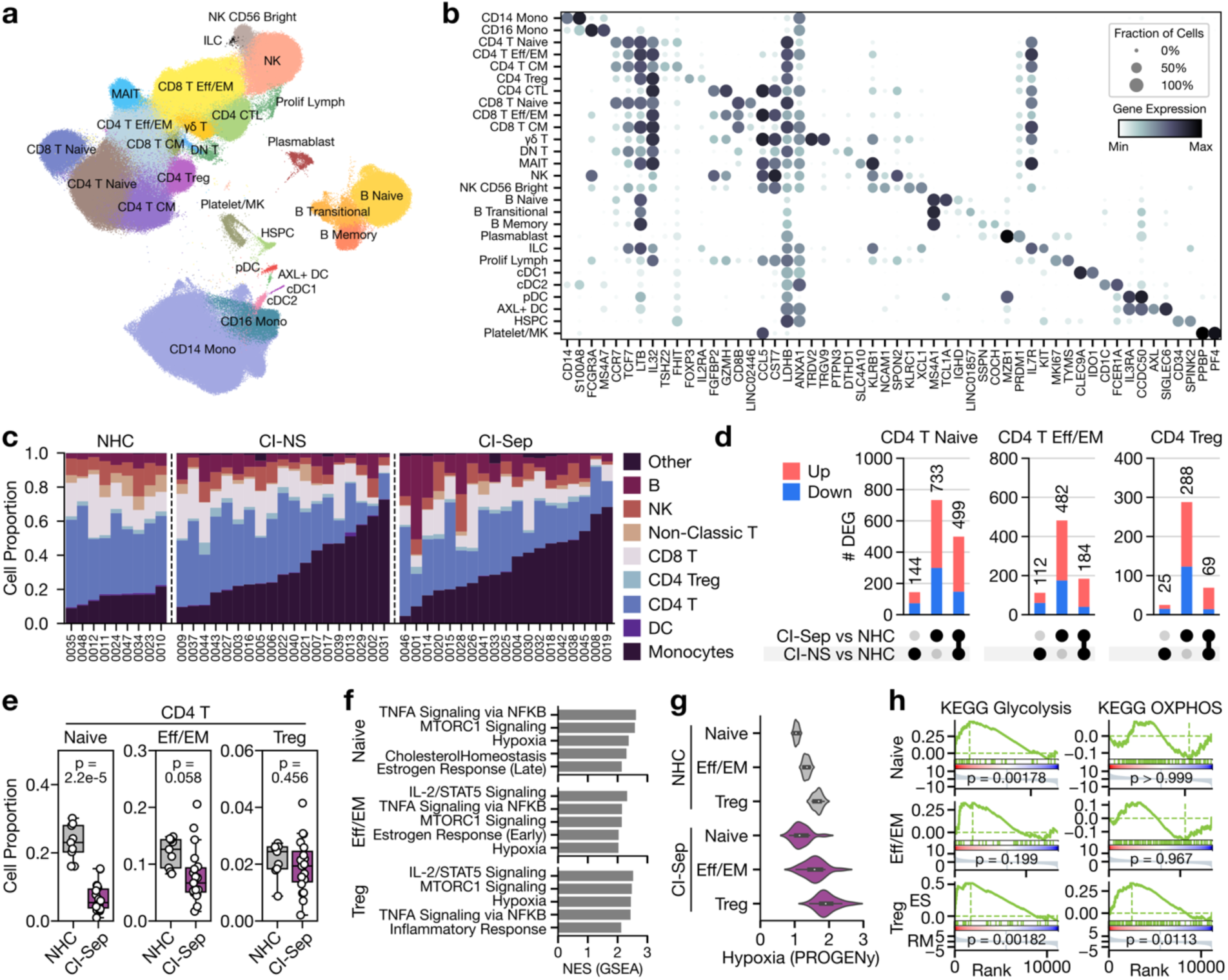
Single-cell transcriptomic map of critical illness and severe sepsis. **a,** UMAP projection of 644,147 cells from 9 NHC, 19 CI-NS, and 19 CI-Sep PBMC samples with detailed cell lineage annotations. **b,** Dot plot summary of canonical cell lineage-defining markers. **c,** Proportionality analysis of major cell lineages with each column representing an individual sample. **d,** Differentially expressed genes (DEG) between CI-NS or CI-Sep and NHC with intersections highlighted as an upset plot. **e,** Proportionality analysis of key CD4 T cell subsets between CI-Sep and NHC. **f,** Top enriched Hallmark pathways by GSEA analysis of pseudobulked scRNA-seq data in CI-Sep compared to NHC for the listed CD4 T cell subsets. **g,** Pathway activity scoring for a consensus hypoxia signature via Pathway RespOnsive GENes (PROGENy). **h,** GSEA assessment of KEGG pathways for glycolysis and oxidative phosphorylation (OXPHOS) in CI-Sep compared to NHC T cell subsets. Statistical analysis for **e** was performed using the Empirical Bayes moderated T-test. Statistical analysis for **h** was performed using the GSEA standard Kolmogorov-Smirnov statistic.

We identified an increased proportion of monocytes with a concurrent reduction in lymphocytes including CD4 T cells consistent with our cytometric data (Fig. 4c). Differential gene expression analysis demonstrated that most transcriptional changes were unique to CI-Sep or shared between CI-NS and CI-Sep (Fig. 4d and Supplemental Table 1). Accordingly, we focused subsequent analyses on CI-Sep versus NHC. Among CD4 T cell subsets, we observed a relative decrease in the proportion of naïve and Eff/EM with preservation of Treg (Fig. 4e). Gene set enrichment analysis (GSEA) using the MSigDB Hallmark gene set collection identified increased TNFα signaling, MTORC1 signaling, IL-2/STAT5 signaling, and hypoxia across CD4 T cell subsets (Fig. 4f). In parallel, we confirmed an elevated hypoxia signature in CI-Sep CD4 T cells using perturbation-informed Pathway RespOnsive GENes (PROGENy) (Fig. 4g). These hypoxia signatures served as evidence that CD4 T cells are subjected to hypoxic stress *in vivo*, likely constraining mitochondrial ATP production and underscoring the critical importance of metabolic plasticity and acquisition of glycolytic capacity, to maintain function under such conditions. Extending GSEA to KEGG metabolic pathways revealed an enrichment of glycolysis in CI-Sep Treg (NES = 1.85, p = 0.00182) and naïve CD4 T cells (NES = 1.67, p = 0.00178) but not in Eff/EM cells (NES = 1.19, p = 0.199) (Fig. 4h). CI-Sep Treg also exhibited a significant enrichment of OXPHOS (NES = 1.53, p = 0.0113), which was unchanged in naïve (NES = -0.63, p > 0.999) and Eff/EM cells (NES = -0.75, p = 0.967. Collectively, these analyses revealed significant changes in cellular composition and transcriptional programming including hypoxia-related signatures and broad, subset-specific metabolic rewiring in CD4 T cells of critically ill septic patients that harmonize with our flow cytometric findings.

### *In Silico* Metabolic Flux Assessment in CD4 T Cell Subsets

To decipher metabolic differences in CD4 T cell subsets from CI-Sep compared to NHC at a more granular level, we applied flux balance analysis (FBA) via Compass^53^. Comparisons of CI- Sep versus NHC CD4 T cell subsets across 22 reaction families revealed disproportionately enriched *in silico* predicted flux in tryptophan metabolism, encompassing kynurenine metabolic pathways (Fig. 5a). Tryptophan, a precursor to serotonin and melatonin, is metabolized by IDO1, IDO2, or TDO2 into N-formyl-L-kynurenine, which is further processed to kynurenine (KYN) (Fig. 5b). KYN can serve as a substrate for *de novo* nicotinamide adenine dinucleotide (NAD) synthesis or as a source of picolinic acid or acetyl-CoA. While CD4 T cells lack IDO1, IDO2, and TDO2, they express SLC7A5 (LAT1), enabling uptake of N-formyl-L-KYN and L-KYN for downstream metabolic processes^54^ . KYN has known immunomodulatory effects on CD4 T cells, including agonism of AHR via KYN metabolites and enhancement of Treg survival and suppressive function^55^. Furthermore, elevated plasma KYN levels are associated with increased morbidity and mortality in sepsis patients^56–60^.

**Figure 5:**
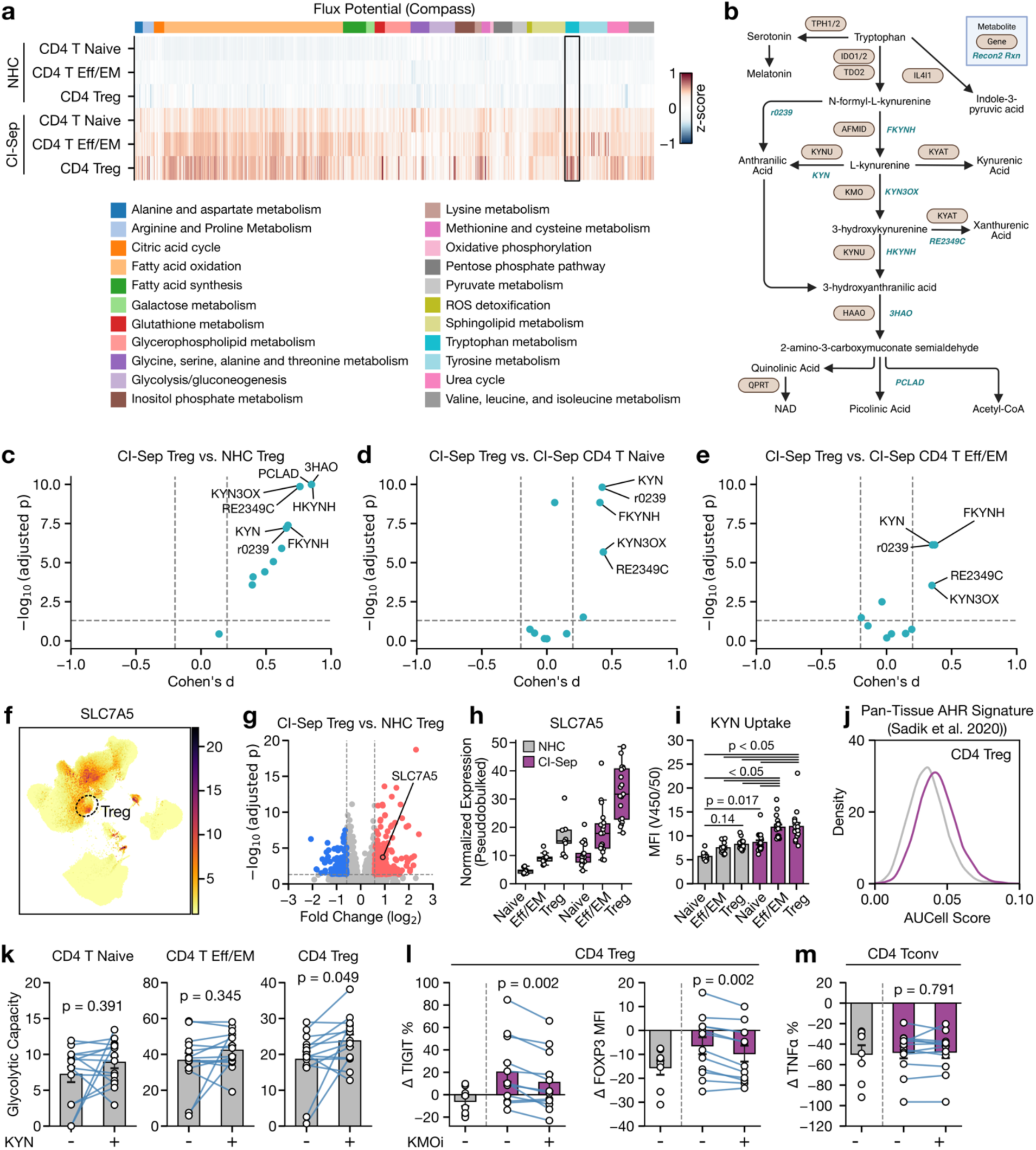
Kynurenine metabolism promotes CD4 Treg glycolytic capacity and suppressive remodeling from critically ill patients with sepsis. **a,** Heatmap visualization of Compass flux balance analysis for the listed major metabolic pathways. Each column represents the reaction score of an individual metabolic reaction from the Recon2 reconstruction scaled via standard z-scoring, with larger values indicating higher predicted flux. Reactions are organized by meta-reaction family as indicated in the legend. **b,** Schematic of tryptophan and kynurenine metabolism with metabolites shown in black, enzymatic genes shown in tan, and associated Recon2 pathways shown in blue. Panel created in BioRender.com. **c,d,e,** Pairwise comparison of effect size of tryptophan/kynurenine metabolism pathways from the Compass analysis in **a**. The magnitude of the difference is represented by the Cohen’s d value (standardized mean difference). **f,** Normalized SLC7A5 transcript expression overlaid on the UMAP representation from Fig 5a. Treg population is outlined. **g,** Differential gene expression analysis by DESeq2 of pseudobulked scRNA-seq data. Genes in red and blue are upregulated and downregulated (>1.5 fold change with adjusted p value < 0.05), respectively, in CI-Sep Treg compared to NHC Treg. **h,** Normalized pseudobulked SLC7A5 transcript expression for the listed CD4 T cell subsets. Each dot represents as individual biologic replicate. **i,** Measured kynurenine (KYN) uptake via fluorescence detection (450 nm filter with 50 nm bandwidth following violet laser excitation) after KYN-pulse. **j,** Pathway activity analysis via AUCell of a consensus human pan-tissue AHR gene signature, with larger values representing higher AHR-associated transcriptional activity, p < 0.0001. **k,** Glycolytic capacity measured by puromycin incorporation assay of paired samples. PBMC were pre-treated for 24 hrs in media alone or media plus 10 µM kynurenine (KYN). **l,** Percentage change in the frequency of TIGIT-expressing Treg and FOXP3 geometric MFI in Treg in anti-CD3/CD28/CD2 restimulated PBMC with stimulation plus oligomycin or stimulation plus oligomycin and KMOi compared to stimulated-only. NHC cohort shown as a reference. **m,** Percentage change in the frequency of TNFα-expressing CD4 Tconv in anti-CD3/CD28/CD2 restimulated PBMC with stimulation plus oligomycin or stimulation plus oligomycin and KMOi compared to stimulated-only. NHC cohort shown as a reference. Statistical analysis for **c,d,e**,**j** was performed using Benjamini-Hochberg (BH)-adjusted Wilcoxon rank-sum testing. Statistical analysis for **g** was performed using the DESeq2 standard BH-adjusted Wald test. Statistical analysis for **i** was performed using the Kruskal-Wallis test followed by Dunn’s post-hoc testing. Statistical analysis for **k-m** was performed using paired sample testing with Wilcoxon signed-rank test.

We focused on pathways associated with tryptophan metabolism from our FBA using pairwise comparisons between CI-Sep Treg and NHC Treg, CI-Sep naïve CD4 T cells, and CI-Sep Eff/EM CD4 T cells. At the reaction level, the most significantly different fluxes were predicted in KYN-specific pathways within tryptophan metabolism across all comparisons (Fig. 5c,d,e). To evaluate the potential for Treg to respond to KYN, we confirmed expression of the KYN transporter SLC7A5 on Treg (Fig. 5f). Differential gene expression analysis demonstrated that SLC7A5 was among the most upregulated genes in CI-Sep Treg compared to NHC Treg (Fig 5g,h).

We next measured the capacity of CD4 T cell subsets to uptake KYN using the natural fluorescence of KYN. PBMC were pulsed with KYN, and intracellular KYN was determined by flow cytometry. KYN uptake was significantly increased in all CI-Sep CD4 T cell subsets compared to NHC, with the highest levels observed in CI-Sep Treg and Eff/EM cells (Fig. 5i and Extended Data Fig. 5a). Consistent with elevated KYN uptake, CI-Sep Treg exhibited an increased transcriptional signature of AHR activity compared to NHC Treg (Fig. 5j)^61^. Collectively, these findings identify differential KYN metabolism as a candidate mechanism for CD4 T cell immunosuppressive remodeling in critically ill patients with sepsis.

### Effect of Kynurenine Metabolism on CD4 T Cell Function

We next investigated the effect of KYN metabolism on CI-Sep CD4 T cell function. PBMCs from NHC were treated with 10 μM KYN for 24 hours followed by glycolytic capacity assessment. This approach utilized a concentration of KYN within the physiologic range measured in the blood of patients with sepsis^56^. This dose was selected to minimize the risk of overestimating the effect size of KYN, a concern that has been raised in prior studies^62^. KYN treatment partially but consistently increased the glycolytic capacity of NHC CD4 Treg, an effect not observed in naïve or Eff/EM CD4 T cells (Fig. 5k). Conversely, to determine if the acquired glycolytic capacity of CI-Sep CD4 T cell subsets could be reversed, we pre-treated CI-Sep PBMC with GSK180, a potent kynurenine-3-monooxygenase inhibitor (KMOi), for 3 hours prior to puromycin incorporation testing. We observed no discernible impact on glycolytic capacity in CD4 naïve, Eff/EM, or Treg subsets (Extended Data Fig. 5b). The neutral impact of KMOi on CI-Sep CD4 T cell subsets was not due to a KYN deficit in our HPLM-based media, which we measured at near physiologic levels (Extended Data Fig. 5c). These findings suggest that KYN contributes to the acquisition of glycolytic capacity in Treg, and this effect is persistent as short-term disruption of KYN metabolism in sepsis-conditioned Treg does not reverse this phenotype.

KYN metabolism promotes suppressive Tregs in healthy subjects^63–65^, though direct experimental evidence for this in systemic inflammatory diseases in humans is sparse. To test whether KYN metabolism contributes the FOXP3 and TIGIT phenotypes in mitochondrially-stressed CI-Sep Treg, we stimulated PBMCs for 24 hours with low-dose oligomycin in the presence or absence of KMO inhibition. KMO inhibition reduced the frequency of TIGIT-expressing Treg and decreased FOXP3 expression, partially reversing the sepsis-associated phenotypes observed in CI-Sep Treg (Fig. 5l). These results highlight a critical role for kynurenine metabolism to maintain Treg vitality and suppressive function in critically ill septic patients.

We also assessed the impact of KMO inhibition on CI-Sep CD4 Tconv. Addition of KMOi reduced the expression of PD-1 and LAG-3, suggesting these CD4 Tconv may be less susceptible to suppression through these immune checkpoint pathways (Extended Data Fig. 5d). KMO inhibition had no significant effect on TNFα- or IFNγ-expressing Tconv, but it led to a modestly greater reduction in the frequency of IL-17A-expressing Tconv compared to oligomycin alone (Fig. 5m and Extended Data Fig. 5e). Additionally, KMOi did not substantially alter the frequency of IL-2-producing Tconv (Extended Data Fig. 5e). Collectively, these findings indicate that disrupting kynurenine metabolism can partially attenuate Treg suppressive phenotypes in critically ill septic patients without major inhibitory effects on Tconv, highlighting a potential target to restore the balance between immunosuppression and traditional CD4 T cell effector functions.

### Treg Glycolytic Capacity Associates with Clinical Metrics and Outcomes

To understand the potential clinical relevance of these metabolic alterations, we assessed the relationship between Treg glycolytic capacity and key ICU clinical metrics and patient outcomes, pooling CI-NS and CI-Sep samples to provide adequate power. Treg glycolytic capacity was positively correlated with key critical illness severity metrics including Sequential Organ Failure Assessment (SOFA) and Acute Physiology and Chronic Health Evaluation II (APACHE II) scores (Fig. 6a,b). No differences in glycolytic capacity were observed by biological sex or age (Fig. 6c,d). While there was no association with peak plasma lactic acid concentration in the first 24 hours, there was a trend with plasma pH, where lower, more acidic pH correlated with higher glycolytic capacity (Fig. 6e,f). Patients with more severe shock denoted by higher vasopressor requirements (norepinephrine equivalents, NEE) exhibited higher Treg glycolytic capacity (Fig. 6g). However, there were no differences in Treg glycolytic capacity when subjects were stratified by alveolar oxygen-exchange abnormalities inferred from supplemental oxygen requirements (SpO2/FiO2 or PaO2/FiO2) (Fig. 6h,i). We previously observed transcriptional signatures of hypoxia in CI-NS and CI-Sep Treg (Fig. 2j and Fig. 4f,g). Given the association of Treg glycolytic capacity with severe shock but not alveolar oxygen-exchange, this suggests that tissue dysoxia rather than arterial oxygen tension may be a more important determinant of Treg metabolism in critical illness.

**Figure 6:**
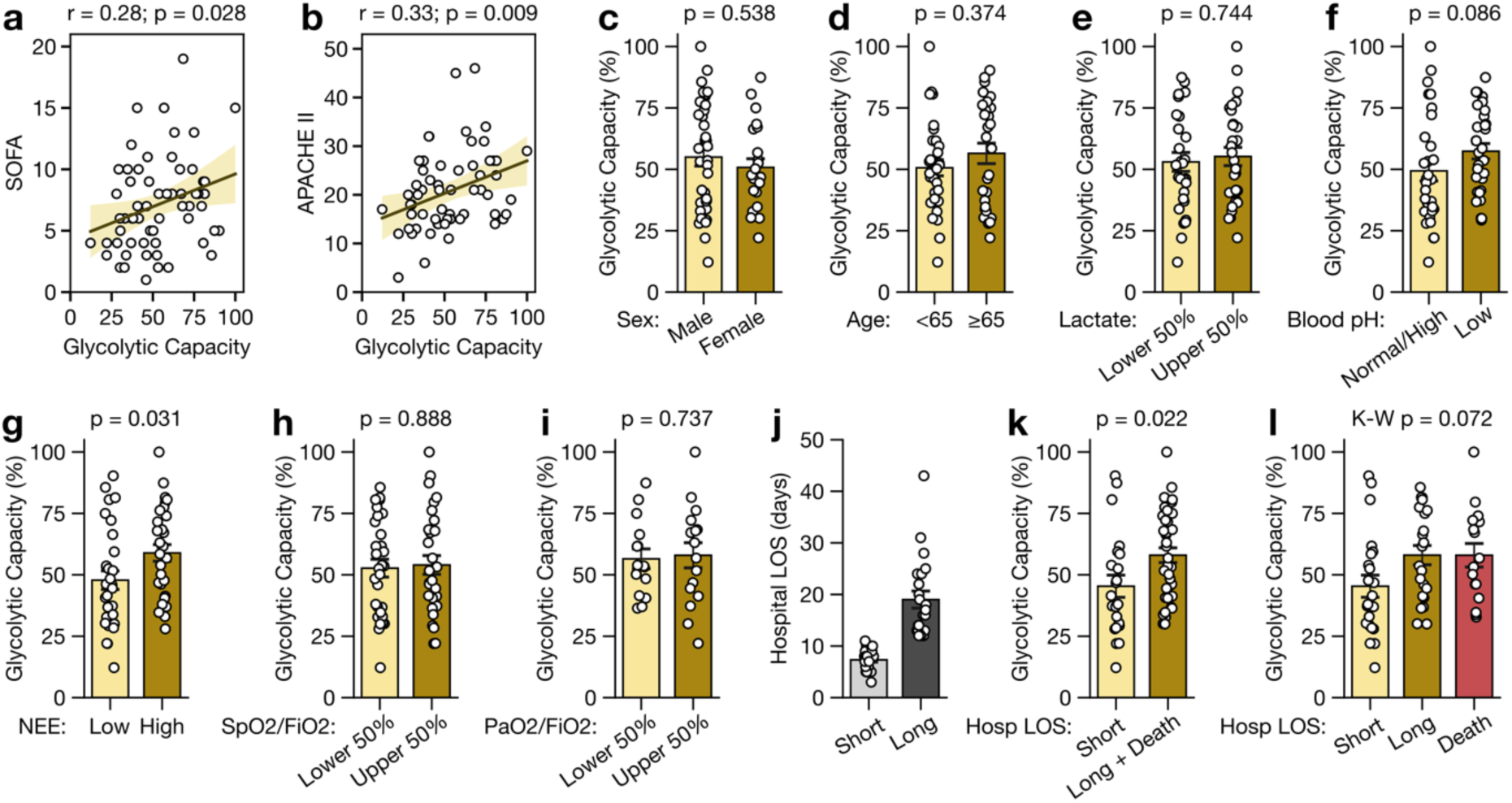
Association of Treg Glycolytic Capacity and Clinical Metrics in Critical Illness. **a,b,** Spearman rank correlation with shaded confidence interval among all critically ill patients (CI-NS + CI-Sep) between Treg glycolytic capacity and SOFA/APACHE II scores. Treg glycolytic capacity stratified by **c,** sex, **d,** age <65 (median = 55 [IQR 47-60]) or ≥65 (median = 71 [IQR 68-78]), **e,** plasma lactate levels into low (lower 50%, median = 1.6 [IQR 1.1-1.8]) and high (upper 50%, median = 4.3 [IQR 3.1-5.8]), **f,** plasma pH into pooled normal/alkalotic (e.g. high) pH (median = 7.40 [IQR 7.37-7.44]) and low pH (median = 7.26 [IQR 7.20-7.29]), **g,** norepinephrine equivalent (NEE) dose into low (lower 50%, median = 0.000 [IQR 0.000-0.036]) and high (upper 50%, median = 0.197 [IQR 0.134-0.389]), **h,** SpO2/FiO2 (S/F) ratio into low (lower 50%, median = 136 [IQR 110-176]) and high (upper 50%, median = 338 [IQR 262-433]), and **i,** PaO2/FiO2 (P/F) ratio into low (lower 50%, median = 124 [IQR 111-146]) and high (upper 50%, median = 338 [IQR 270-382]). **j,** Hospital length of stay (LOS) in days among survivors to hospital discharge stratified into the shortest 50% (median = 7 [IQR 6-8]) and longest 50% (median = 16 [IQR 13-23]). Arterial blood gas measurement of PaO2 in **s** was not performed by the treating clinical team in all patients; values shown where available. Statistical analysis for **c-i,k** was performed using the Mann-U Whitney test. Statistical analysis for **l** was performed using the Kruskal-Wallis test with Dunn’s post-hoc testing. Error bars represent SEM. K-W = Kruskal-Wallis.

Primary illness severity and the acquisition of secondary complications can prolong hospital stays or lead to in-hospital mortality. We dichotomized survivors to hospital discharge into the shorter or longer 50% of hospital length of stays (LOS), with median LOS 7 and 16 days, respectively (Fig. 6j). Patients with longer hospital LOS or in-hospital death had significantly higher Treg glycolytic capacity compared to those with shorter LOS, with similar effects observed when long LOS and in-hospital death were separated (Fig. 6k,l). Collectively, these findings highlight a strong association between metabolic remodeling in Treg and worse clinical metrics including higher SOFA and APACHE II scores as well as poor clinical outcomes including more severe acidemia, worse shock, prolonged hospital LOS, and mortality.

## DISCUSSION

The immunologic remodeling toward pathologic immunosuppression with poor clinical outcomes is well-documented in critical illness with and without sepsis, yet the molecular mechanisms driving this shift in humans remain poorly understood. Here, we demonstrate that CD4 T cells in critical illness undergo subset-specific metabolic remodeling, favoring the persistence of suppressive Treg and the attrition of Tconv. Of CD4 T cell subsets, Treg displayed the greatest metabolic plasticity, acquiring substantial glycolytic capacity relative to naïve and Eff/EM CD4 T cells, correlating with their preserved frequency in critical illness. These findings provide evidence of subset-specific metabolic reprogramming among CD4 T cells in human critical illness and sepsis, emphasizing the importance of analyzing individual T cell subsets to determine mechanisms of immune dysfunction. Additionally, sepsis metabolic studies in humans have often focused on dysfunctional pathways, such as impaired mitochondrial function^38,66^. Our findings highlight that newly acquired metabolic capacities may also play critical roles, with the integrated balance between these losses and acquisitions contributing to immunologic remodeling in sepsis.

The magnitude of Treg metabolic remodeling correlates with worse clinical metrics and outcomes, including elevated SOFA and APACHE II scores, more severe shock, and prolonged hospital stays or death. While our framework does not allow us to definitively establish causality of this association, our overall results demonstrate mechanistically that this remodeling favors Treg survival and suppressive function over conventional T cells—a shift broadly implicated in patient morbidity and mortality^9,17^.

Our findings underscore the importance of accounting for disease-specific conditions such as mitochondrial stress, which is pervasive in sepsis and critical illness and evident in our analyses. Functional assessments in the presence of mitochondrial impairment revealed unique Treg properties in critically ill and septic patients, including FOXP3 stabilization and TIGIT upregulation. The coexistence of cellular and metabolic stressors may also reconcile prior discordant data on the magnitude of KYN-induced immunosuppression^62^, which was unmasked in our studies in the presence of mitochondrial disruption.

We identify KYN metabolism as a key driver of immunometabolic remodeling in septic ICU patients. Kynurenine promoted Treg-centric metabolic shifts in CI-Sep, including enhanced glycolytic capacity and stabilization of immunosuppressive markers such as FOXP3 and TIGIT. While its effects were partial—reflecting the complex physiologic changes in sepsis—our findings harmonize with prior clinical data linking elevated plasma kynurenine levels with poor patient outcomes, including mortality^56–60^. Moreover, we provide evidence for kynurenine’s direct role in modulating CD4 T cells in a human inflammatory disease context, expanding on prior studies focused on healthy human T cells^63–65^. Tryptophan and kynurenine metabolism represent highly druggable targets with translational potential. IDO1 and TDO inhibitors are under active investigation in cancer^67^, where they aim to mitigate tumor immune evasion, while KMO inhibitors are being developed for neurodegenerative diseases and pancreatitis^68,69^. These agents, which have demonstrated safety in clinical trials, may represent candidate therapeutic approaches to recalibrate immune function in critical illness and sepsis. Importantly, the effects of KMO inhibition in our *ex vivo* studies were partial, which may represent a potential advantage in this therapeutic approach. Sepsis is an inherently heterogeneous condition, characterized by significant variability at both the patient and pathogen levels including in the degree of hyperinflammatory and immunosuppressive dysfunction. In our analyses, KMO inhibition attenuated sepsis-associated Treg phenotypes without entirely abolishing Treg function or inducing Tconv hyperactivation. This precision-based strategy aims for balanced restoration of immune homeostasis that could mitigate pathologic immunosuppression while avoiding the risks of excessive effector activation.

In contrast to other immunomodulatory strategies, tryptophan and kynurenine metabolism operates across both immune and non-immune tissues, presenting broader therapeutic potential. For example, kynurenine modulates vascular tone as an arterial vasodilator^70^, and thus disruption of kynurenine metabolism may improve blood pressure and perfusion in septic shock. Additionally, KMO inhibition shifts kynurenine metabolism toward neuroprotective metabolites like kynurenic acid while reducing neurotoxic derivatives such as 3-hydroxykynurenine and quinolinic acid^71–73^. This is particularly relevant as elevated plasma KYN levels have been associated with prolonged delirium in ICU patients yet there are no pharmacologic interventions to treat critical illness-associated brain dysfunction^74,75^. Other kynurenine metabolites, notably 3-hydroxykynurenine, have been shown to be cellular and tissue toxins that may perpetuate end-organ injury in critical illness^76–80^. Preclinical models support the potential of KMO inhibition to mitigate this end-organ injury, including acute kidney injury in a mouse model of ischemia/reperfusion^81^ and multiorgan injury in rodent models of acute pancreatitis^80,82^. Collectively, these findings suggest that targeting kynurenine metabolism could provide a multifaceted therapeutic approach, addressing not only immune dysfunction but also broader pathophysiologic consequences of critical illness.

This study has several limitations. First, while our ICU cohort included medical and surgical patients, the generalizability of these findings to trauma or burn populations where sepsis is also common is unclear. Second, most of our analyses were performed using samples collected at day 2 post-ICU admission, limiting our ability to disentangle longitudinal complexities. Where studies were performed with serial sampling through day 7, factors such as clinical improvement leading to discharge and patient deaths provide some uncertainty to our assessments as day 7 samples may not fully represent the enrollment cohort. In addition, our focus on subsets of CD4 T cells, including rare subsets like Treg, was conducted in the context of critical illness-associated lymphopenia and limited blood volumes due to patient severity of illness. These factors restricted our ability to perform certain metabolic assays in Treg, such as heavy isotope tracing and extracellular flux analysis, that require large numbers of cells. Attempts to expand Treg via standard protocols abolished the sepsis-associated metabolic phenotypes observed by puromycin incorporation assay and were thus not exploitable. Finally, our clinical correlations with Treg glycolytic capacity were underpowered when stratified by sepsis versus non-sepsis, leaving the sepsis-specificity of these findings uncertain.

Despite these constraints, our study uniquely includes a diverse array of non-septic ICU controls, exclusively utilizes human biospecimens, and represents the largest published PBMC scRNA-seq dataset to date in non-COVID critical illness. Our focus on sepsis in the critical care setting, specifically among the most severely ill patients (median SOFA 8 in CI-Sep), complements prior large sepsis scRNA-seq studies that included broader cohorts encompassing both ICU and non-ICU patients (e.g., median SOFA 2-4^83^ and 5^84^). While changes in CI-Sep generally exhibited larger effect sizes, the directionality of immunometabolic changes was consistent between CI-NS and CI-Sep, suggesting overlapping mechanisms of metabolic dysfunction in severe critical illness. However, transcriptional analysis revealed numerous genes unique to CI-Sep, indicating that meaningful differences may still exist between CD4 subsets in CI-Sep and CI-NS that warrant further investigation. Together, we identify kynurenine metabolism as a key modulator of CD4 T cell immunometabolic rewiring in critically ill patients with sepsis supporting the persistence of suppressive Tregs at the expense of Tconv. These findings underscore the potential of targeting kynurenine metabolism to restore immune homeostasis and improve clinical outcomes. This work highlights the importance of precision-based approaches to addressing immune dysfunction in critically ill patients.

## AUTHOR CONTRIBUTIONS

M.T.S. and J.C.R. designed the research and prepared the manuscript with contributions from all other authors. M.T.S., A.E.S., C.M.N., and E.Q.J. performed the research and analyzed data. P.L. provided bioinformatics technical support. E.M.W. provided technical support with methods development. D.C.N provided essential intellectual support in experimental design and planning. J.A.B and L.B.W. provided essential expertise and clinical samples. H.H.D. provided key technical support. E.L.M, C.Y.S, S.M.T., C.M.N, M.T.S., and A.E.S. collected and/or processed clinical samples.

## METHODS

### Human Subjects, Clinical Metrics, and Sepsis Adjudication

Critically ill patients were recruited from the medical and surgical intensive care units (ICUs) at Vanderbilt University Medical Center (VUMC) from 2022 to 2024 as part of The Sepsis ClinicAl Resource And Biorepository (SCARAB). Patients reported in this study were included if they were ≥18 years of age and did not meet any of the study exclusion criteria: active exsanguination without hemostatic control or a current hemoglobin ≤7.5 g/dL, ICU stay at VUMC or an outside facility of >48 hours, cardiac arrest prior to enrollment, moribund with imminent death, receiving extracorporeal membrane oxygenation (ECMO) at the time of enrollment, pregnant, medical trainees, in police custody, admitted to the ICU solely for frequency of nursing care, or admitted for an uncomplicated diabetic ketoacidosis, drug overdose, or alcohol withdrawal. For patients reported in this manuscript, we additionally excluded any patients who were receiving T cell directed immunosuppression (e.g. tacrolimus, azathioprine, mycophenolate, etc.), receiving immunomodulatory antineoplastic agents (e.g. pembrolizumab, ipilimumab, rituximab, etc.), had severe leukopenia with insufficient PBMC for analysis, had an active hematologic malignancy, or had received an allogenic stem cell transplant. Informed consent was obtained from patients or their surrogates. If informed consent could not be obtained due to lack of patient capacity or inability to reach a listed surrogate, then patients were enrolled in this minimal risk study under an Institutional Review Board (IRB)-approved waiver of consent. Community-matched non-acutely ill healthy controls were recruited from bedside surrogates of enrolled ICU patients. Participants were required to be ≥18 years of age and not meeting an exclusion criteria: pregnant, presence of an active acute infection, medical history of significant anemia, receiving T cell directed immunosuppression, currently receiving any antineoplastic agents, presence of an active hematologic malignancy, or previously receiving an allogenic stem cell transplant. Written informed consent was obtained for all heathy controls. These studies were approved by the IRB at VUMC under protocols #211462 (SCARAB) and #191562 (healthy controls).

For ICU patients, peripheral blood was obtained within 24 hours of ICU admission (day 0) and then serially 2 and 7 days after study enrollment. Unless otherwise stated, all experiments were performed with samples collected on day 2. For PBMC isolation, a total of approximately 7, 11, and 11 mL blood were obtained at each respective timepoint in a combination of citrate and EDTA vacutainer tubes. The indication for collecting both citrate and EDTA tubes pertained to lines of investigation unrelated to this current study; to maximize the small volume of blood we obtained per patient, we chose to pool PBMC from these two vacutainer types. Clinical data, including demographics, medical history, vital signs, laboratory data, imaging reports, medications, and flowsheet data, was recorded from the chart and stored in a HIPAA-compliant REDCap database. For healthy control participants, up to 50 mL of blood was obtained at a single timepoint in a combination of citrate and EDTA vacutainer tubes. Clinical data for healthy control participants was limited to age and biologic sex, which were recorded in a HIPAA-compliant REDCap database.

Sepsis status among critically ill patients was adjudicated post-hospitalization by a practicing physician intensivist (Extended Data Fig. 1). The presence of an infection was established based on positive bacterial cultures, infection-equivalent findings such as a visibly contaminated abdomen during laparotomy, or through integration of the totality of clinical data in the absence of the former. For patients with confirmed or highly suspected infections, evidence of end-organ injury was assessed using Sepsis-3 criteria, defined as a sequential organ failure assessment (SOFA) score increase of ≥2 from the patient’s baseline^2^. Patients with established infections and evidence of end-organ injury due to the infection were adjudicated as having sepsis. Critically ill patients who did not meet sepsis criteria were included in the non-sepsis group.

### Peripheral Blood Mononuclear Cell (PBMC) Isolation and Storage

Whole blood was pooled from EDTA and citrated tubes, diluted 1:2 with PBS, and layered over 15 mL of Lymphoprep density gradient medium (STEMCELL Tech., Cat. #18061) in a SepMate-50 (IVD) tube (STEMCELL Tech., Cat. #85450). Density gradient preparations were then centrifuged at 1200g for 15 minutes at 20°C and the top layer containing PBMC and plasma was poured into a new 50 mL conical tube. The PBMC/plasma mixture was then spun at 400g for 10 minutes at 20°C and the supernatant was aspirated off. The cell pellet was resuspended in 4 mL ACK lysing buffer (Gibco, Cat. #A1049201) for 7 minutes and then 36 mL of PBS was added to quench the lysing reaction. This mixture was then spun at 300g for 10 minutes at 20°C and the supernatant aspirated off. The cells were then resuspended in 30 mL PBS, filtered through a 70 µm strainer into a new 50 mL conical tube, and then spun at 300g for 10 minutes at 20°C. The supernatant was aspirated off and the cells resuspended in Bambanker Cell Freezing Medium (GC LYMPHOTEC Inc., Cat. #BB02) at a concentration of 2 million PBMC per mL. PBMC were aliquoted and slow frozen in a CoolCell FTS30 freezing container (Corning, Cat. #432006) at -80°C. Cells were transferred to liquid nitrogen vapor phase storage within 72 hours.

### Cryorecovery of PBMC

PBMC were thawed from cryopreservation by rapid rewarming at 37°C for 3 minutes. Cell suspensions were diluted 1:5 in Human Plasma-Like Medium (HPLM) (Gibco, Cat. #A4899101) supplemented with 10% dialyzed fetal bovine serum (dFBS) (Sigma-Aldrich, Cat. #F0392) and 100 units/mL of penicillin plus 100 μg/mL of streptomycin (P/S) (Gibco, Cat. # 15140122). Diluted cell suspensions were centrifuged at 300g for 5 minutes at 20°C and the supernatant was aspirated off. Two additional washes were performed in HPLM with 10% dFBS and P/S. The final centrifugation step was performed using a 5 mL round-bottom polystyrene test tube with a 35 μm cell strainer cap to filter any remaining debris. After aspiration of the supernatant, the cells were resuspended in HPLM with 10% dFBS and P/S and used in downstream experiments.

### Flow Cytometry

PBMC cell suspensions, either direct from cryorecovery or after incubation, were distributed in 96-well V-bottom microplates (Corning, Cat. #3894). Cells were centrifuged at 300g for 5 minutes at 4°C and the supernatant was removed. Cells were similarly washed twice in cold PBS. Washed cells were resuspended in 50 μL PBS with Ghose Dye Red 780 Fixable Viability Dye (Cell Signaling, Cat. #18452) diluted 1:2000 and Human TruStain FcX solution for Fc receptor blocking (BioLegend, Cat. #422302) diluted 1:40 and incubated for 30 minutes at 4°C. Cells were washed with flow staining buffer (cold PBS + 2% FBS), resuspended in 50 μL of surface antibody stains (antibodies in Extended Data Table 2 diluted in flow staining buffer), and incubated for 30 minutes at 4°C. Cells were twice washed with flow staining buffer. In some experiments, intracellular markers were evaluated. To measure these, washed cells were resuspended in 100 μL of Fix/Perm reagent (Tonbo Biosciences, Cat. #TNB-0607-KIT) and incubated for 30 minutes at room temperature. Subsequently, fixed cells were washed twice with permeabilization buffer (Tonbo Biosciences, Cat. #TNB-1213-L150), resuspended in 50 μL of intracellular antibody stains (antibodies in Extended Data Table 2 diluted in permeabilization buffer), and incubated for 30 minutes at 4°C. Cells were washed twice in permeabilization buffer, resuspended in flow staining buffer, and acquired on a Miltenyi MACSQuant Analyzer 16 flow cytometer. Data exported as MQD files were analyzed in FlowJo Version 10.9.0 (Becton Dickson & Company). Doublets were removed based on forward/side scatter properties and dead cells (viability dye positive) were excluded in all analyses. Representative gating strategies are shown in Extended Data Fig. 6 and full gating strategies for all cell types assessed are listed in Extended Data Table 3.

### Puromycin Incorporation Assay

We measured metabolic dependencies and capacities using a flow cytometric puromycin incorporation assay. Our protocol is adapted from the initial iteration of this method, which was reported as Single Cell ENergetIc metabolism by profilIng Translation inHibition (SCENITH)^45^. After cryorecovery, cells from each sample were split into the indicated number of wells of a 96 well plate and incubated in HPLM with 10% dFBS and P/S for 3 hours at 37°C to allow for restoration of their metabolic program after frozen storage. For each sample, one metabolic inhibitor was added per well; an additional well had no metabolic inhibitors and the final well had a pool of all the inhibitors to fully extinguish ATP production. These were incubated for 30 minutes at 37°C. The following inhibitors were used: glucose-dependent pathway inhibitor 2-deoxy-D-Glucose (2-DG) at a final concentration of 100 mM (Cayman Chemical, Cat. #14325), mitochondrial ATP synthase inhibitor Oligomycin A at a final concentration of 1.5 μM (Cayman Chemical, Cat. #11342), or glutaminase 1 (GLS1) glutaminolysis inhibitor CB-839 at a final concentration of 10 μM (MedChemExpress, Cat. #HY-12248), and carnitine palmitoyltransferase-1 (CPT-1) fatty acid oxidation inhibitor Etomoxir at a final concentration of 5 μM (Sigma-Aldrich, Cat. #E1905). Cells were then pulsed with puromycin at a final concentration of 10 μM (Sigma-Aldrich, Cat. #P8833) and incubated for an additional 40 minutes at 37°C. Cells were then washed in cold PBS and further prepared for flow cytometry as described above. To measure puromycin incorporation, we stained fixed and permeabilized cells using an Alexa Fluor 488-tagged anti-puromycin antibody (Millipore-Sigma, clone 12D10 at 1:1000 dilution, Cat. #MABE343-AF488). Metabolic dependencies and capacities were calculated from the geometric MFI of puromycin within each cell population as originally reported^45^. In some experiments, cells were treated for 24 hours at 37°C in HPLM with 10% dFBS and P/S with 10 µM L-kynurenine (MedChemExpress, Cat. #HY-104026) prior to the puromycin incorporation assay. In some experiments, cells were treated for 3 hours at 37°C in HPLM with 10% dFBS and P/S with 10 µM GSK180 (MedChemExpress, Cat. #HY-112179), a kynurenine-3-monooxygenase (KMO) inhibitor, prior to the puromycin incorporation assay.

### Mitochondrial Mass, Membrane Potential, and Reactive Oxygen Species (ROS)

Mitochondrial mass was determined using MitoTracker Green FM (Invitrogen, Cat. #M46750) and mitochondrial membrane potential was determined using Tetramethylrhodamine Ethyl Ester (TMRE, Invitrogen, Cat. #T669). Cryorecovered PBMC were resuspended in HPLM with 10% dFBS and P/S and plated in a 96 well plate at 37°C for 30 minutes to allow for mitochondrial recovery from a frozen state. PBMC were then treated with MitoTracker Green FM (final concentration 25 nM) and TMRE (final concentration 40 nM) and incubated for an additional 30 minutes at 37°C. Cells were then washed twice with cold flow staining buffer (PBS + 2% FBS) and a final time with cold PBS. Cells were then processed for flow cytometry including viability dye, Fc block, and surface marker antibodies and analyzed as described above. Mitochondrial ROS were determined using MitoSox Green superoxide indicator (Invitrogen, Cat. #M36006). Cryorecovered PBMC in HPLM with 10% dFBS and P/S rewarmed for 30 minutes at 37°C were treated with MitoSox Green (final concentration 500 nM) and incubated for an additional for 30 minutes at 37°C. Cells were subsequently washed and processed for flow cytometry as described above.

### *Ex vivo* Restimulation for CD4 T Cell Function

Cryorecovered PBMC were resuspended in HPLM with 10% dFBS and P/S and plated in a 96 well plate. T cells were stimulated with a 1:20 dilution of ImmunoCult Human CD3/CD28/CD2 T Cell Activator (STEMCELL Tech., Cat. #10970). In some wells, 10 nM oligomycin (sublethal dose) was added at the time of plating and stimulus as a source of mitochondrial stress. Cells were incubated for 24 hours at 37°C. At 5 hours prior to the experimental endpoint, the protein transport inhibitor Brefeldin A (GolgiPlug, BD, Cat. #555029) was added at 1:1000 dilution to allow for intracellular cytokine detection. At 24 hours, cells were collected and analyzed by flow cytometry. In some experiments, some cells were also treated at the time of plating with 10 µM GSK180 (MedChemExpress, Cat. #HY-112179), a kynurenine-3-monooxygenase inhibitor, for the duration of the stimulation.

### Kynurenine Uptake Assay

Intracellular kynurenine uptake was assessed as previously described^54^ with modifications. Cryorecovered PBMC were resuspended in HPLM with 10% dFBS and P/S and plated in a 96 well plate. As a control in some experiments, 10 mM BCH (MedChemExpress, Cat. #HY-108540) was added at the time of PBMC plating to inhibit the transporter activity of SLC7A5. Cells were incubated at 37°C for 1 hour. Cells were washed twice with warm PBS and incubated with Fc receptor blocker, viability dye, and cell surface antibodies diluted in PBS at room temperature for 30 minutes. Cells were washed once with HBSS warmed at 37°C and resuspended in 37°C HBSS with 100 µM L-kynurenine (MedChemExpress, Cat. #HY-104026). In some instances, 10 mM BCH was continued at this step. As an additional control of receptor specificity, a saturating dose of the SLC7A5 substrate L-leucine at 5 mM (MedChemExpress, Cat. #HY-N0486) was added at the time of L-kynurenine administration. As a control for uptake versus cell surface binding, in some instances L-kynurenine chilled at 4°C was added and cells were incubated on ice. These incubations with L-kynurenine were performed for 5 minutes at 37°C followed by reaction quenching with 100 µL IC Fixation Buffer (Invitrogen, Cat. #00-8333) for 30 minutes at room temperature. Samples were washed twice in PBS + 2% FBS and analyzed by flow cytometry. L-kynurenine has spectral properties that can be excited by a 405 nm violet laser and emission captured on a 450/50 nm detector (V1 channel on the MACSQuant Analyzer 16).

### LC-MS/MS Quantitation of Kynurenine in Cell Culture Media

To determine the pre-existing availability of L-kynurenine in the media preparations used in experiments where modulations included adding exogenous L-kynurenine or a kynurenine-3-monooxygenase inhibitor (KMOi), we performed quantitative liquid chromatography with tandem mass spectrometry. L-kynurenine levels were tested in fresh HPLM, fresh HPLM plus 10% dFBS and P/S, and HPLM plus 10% dFBS and P/S stored for 30 days at 4°C. 1 mL of -80°C 80:20 MeOH:H_2_O was added to the 330 µL of media sample in a 1.7 mL tube and vortexed vigorously. Then, 20 nmol of internal standard (^13^C-1-Lactate, MilliporeSigma, Burlington, MA) was spiked into each sample. Cells were then placed in -80°C freezer to extract for 15 minutes. After extraction, any insoluble debris was pelleted at 16,000 x g for 10 minutes at 4°C. The supernatant containing extracted metabolites was then transferred to a new 1.7 mL tube. Extracts were then dried under N_2_. Dried samples were resuspended in 50 µL 3:2 mobile buffer A: mobile buffer B (see below) and transferred to 1.7 mL tubes. Samples were then centrifuged at 16,000 x g for 10 minutes at 4°C to remove any insoluble debris. A standard curve was generated using 20 nmol of internal standard (^13^C-1-Lactate) in samples containing either 0, 3.75, 7.5, 15, or 30 nmol of a natural isotope kynurenine standard. 18 µL of the sample was then chromatographed with a Shimadzu LC system equipped with a 100 x 2.1mm, 3.5μm particle diameter XBridge Amide column (Waters, Milford, MA). Mobile phase A: 20 mM NH_4_OAc, 20 mM NH_4_OH, 5% acetonitrile in H_2_O, pH 9.45 (pH with NH_4_OH); mobile phase B: 100% acetonitrile. With a flow rate of 0.45 mL/min the following gradient was used: 2.0 min, 95% B; 3.0 min, 85% B; 5.0 min, 85% B; 6.0 min, 80% B; 8.0 min, 80% B; 9.0 min, 75% B; 10 min, 75% B; 11 min, 70% B; 12 min, 70% B; 13 min, 50% B; 15 min, 50% B; 16 min 0% B; 17.5 min, 0% B; 18 min, 95% B. The column was equilibrated for 3 minutes at 95% B between each sample. Scheduled MRM was conducted in negative and positive mode with a detection window of 120 seconds using an AB SCIEX 6500 QTRAP with the following analyte parameters: *m/*z 209 → 94 (positive mode, retention time: 6.3 minutes) for L-kynurenine; *m/z* 90 → 43 (negative mode, retention time: 6.0 minutes) for ^13^C-1-lactate internal standard. All analytes were quantified via LC-MS/MS using the standard curve.

### Single-Cell Library Preparation and RNA Sequencing

Cryopreserved PBMCs (0.5–1.0 mL, 2 million cells/mL) were rapidly thawed at 37°C for 3 minutes, then diluted dropwise into 7 mL of HPLM supplemented with 10% dialyzed fetal bovine serum (dFBS). Cells were centrifuged at 300 × g for 5 minutes at room temperature, washed with 2 mL of HPLM + 10% dFBS, and subsequently washed with 2 mL of cold PBS containing 0.04% bovine serum albumin (BSA). The final wash step included filtration through a 35-μm cell strainer (Falcon, Cat. #352235) during centrifugation. Cells for single-cell RNA-seq library generation were resuspended in cold PBS + 0.04% BSA at a concentration of 1,000 cells/μL. Single-cell RNA-seq libraries were generated using the Chromium Next GEM Single Cell 5’ HT Kit v2 (10x Genomics) targeting 20,000 cells per sample for capture. Libraries were sequenced on an Illumina NovaSeq 6000 system (S4 flow cell, PE150) targeting 50,000 reads per cell. Raw sequencing data were processed using Cell Ranger software (v7.1.0), aligning reads to the GRCh38-2020-A transcriptome reference with the intron inclusion option enabled.

### scRNA-Seq Preprocessing, Quality Control, Integration, and Modeling

scRNA-seq processing was conducted using a custom generated Nextflow workflow via AWS HealthOmics. Raw feature barcode matrix outputs from Cell Ranger were stored in an AWS S3 bucket and used as the initial input. Ambient RNA and technical artifacts were removed using the remove-background function of CellBender (v0.3.0)^85^ executed within a Docker container built from the base image nvidia/cuda:11.7.1-runtime-ubuntu20.04 with Python (v3.7). Learning rates (0.0000005-0.00005) and training epochs (150-300) were iteratively tuned, while all other settings were kept at their default values.

Outputs from CellBender were filtered using Seurat (v4.3.0)^86^ to exclude cells with mitochondrial content >20% and feature counts <400 or >5000. Doublets were removed using scDblFinder (v1.13.10)^87^, informed by temporary normalization and variance stabilization with SCTransform and shared nearest neighbor (SNN) graph clustering (resolution=0.5) based on the top 2,000 highly variable genes (HVGs) and 30 principal components. This normalization and clustering was used exclusively for doublet detection and discarded thereafter. Sample-level metadata was added to each file, and the cleaned dataset of high-quality singlet cells was converted into an AnnData object using a custom implementation of the sceasy package (v0.0.7)^88^. The final AnnData object retained the filtered raw count matrix and sample-, cell-, and gene-level metadata. These steps were executed within a Docker container built from the base image bioconductor/bioconductor_docker:RELEASE_3_17 with R (v4.3.0) and Python (v3.9) accessed via the R reticulate (v1.28) interface. Of note, due to grossly failing quality control metrics, one NHC sample (0036) was excluded from the analysis, leaving 9 high-quality NHC samples for downstream processing.

Post-quality control, single-cell RNA-seq data were integrated and modeled using scVI-tools (v1.0.4)^89^. Individual AnnData files were merged into a combined object, normalized (target_sum=1e4), and log-transformed using Scanpy (v1.9.5)^90^. The top 5,000 HVGs were identified using highly_variable_genes (flavor=’seurat_v3’). The scVI model was trained with n_hidden=128, n_latent=30, and n_layers=2, incorporating categorical covariates (Biologic_Sex) and continuous covariates (percent.mt, nCount_RNA, and percent.ribosomal), and accounting for batch effects (batch_key). A negative binomial likelihood (gene_likelihood = ’nb’) with gene-level dispersion and a dropout rate of 0.1 was used to model gene expression. The model was trained for a maximum of 400 epochs with early stopping, triggered after 50 epochs without improvement in validation ELBO loss. Validation was performed every epoch, and the AdamW optimizer was used with a learning rate of 0.001. Denoised HVG expression values were generated using the scVI get_normalized_expression function (library_size=1e4, n_samples=25) and stored as a separate layer in the AnnData object. These steps were executed in a Docker container built from the base image nvidia/cuda:11.7.1-runtime-ubuntu20.04 with Python (v3.9), PyTorch (CUDA 11.7 optimized), and JAX (CUDA 11 support).

### scRNA-seq Clustering, Visualization, and Annotation

Scanpy (v1.9.5) was used to create a k-nearest neighbors (kNN) graph (scanpy.pp.neighbors) using the scVI latent space (use_rep=’X_scVI’, n_neighbors=30). Initial clustering was conducted with the Leiden algorithm (scanpy.tl.leiden, resolution=1). The most differentially expressed genes for each cluster were identified via the differential_expression function using the scVI model with denoised expression. Cell type annotations were initially assigned based on canonical marker gene expression, using the Azimuth Human PBMC core reference map as the primary guide^91^, supplemented by insights from published literature and expert consensus. Ambiguous clusters without dominant marker profiles were iteratively subclustered (resolutions between 0.2–2.0) until high-confidence annotations could be assigned. Clusters exhibiting two distinct, non-overlapping gene expression profiles were excluded as potential doublets. A small B cell population in sample V-0072.2 expressing canonical markers of chronic lymphocytic leukemia (ROR1, FMOD, CD5) and exhibiting identical heavy and light chains was excluded as a likely subclinical neoplasm. Final cell type annotations were provided at two levels: celltype_coarse for major lineages and celltype_fine for granular subpopulations. Annotated clusters were visualized using Uniform Manifold Approximation and Projection (UMAP) via the scanpy.tl.umap and scanpy.pl.umap functions. A dot plot was generated using scanpy.pl.dotplot to visualize the expression of manually selected cell lineage marker genes.

### Cell Proportionality

To determine the proportion of each cell type within the sample, we used the Scanpro method^92^. Scanpro was run on NHC, CI-NS, and/or CI-Sep samples with both celltype_coarse and celltype_fine cell lineage annotations. Statistical comparisons were made using the Empirical Bayes moderated T-test for pairwise comparison, adjusted for multiple testing with the Benjamini-Hochberg method.

### Differential Gene Expression and Pathway Analysis

Pseudobulk differential gene expression analysis was conducted using the python implementation of the decoupleR package^93^. Aggregated pseudobulk profiles were generated from single-cell data by summing counts for each cell type across samples, retaining profiles with at least 20 cells and 1,000 total counts per pseudobulk. Differential gene expression was analyzed for CD4 T cell subsets across all group (NHC, CI-NS, and SI-Sep) comparisons using PyDESeq2^94^. Genes with very low or sporadic expression within a group were excluded (filter_by_expr function, min_count=3, min_total_count=25). Log2 fold change and false discovery rate (FDR)-adjusted p-values were calculated via PyDESeq2.

Gene Set Enrichment Analysis (GSEA)^95^ was performed on pseudobulk differential gene expression results using decoupleR. CD4 T cell subsets were compared between CI-Sep and NHC, with the Wald statistic from DESeq2 serving as the ranking metric. We evaluated the Hallmark gene set collection and selected KEGG gene sets from MSigDB. Enrichment scores were calculated using 1,000 permutations, considering gene sets with at least 15 genes. A random seed (42) was set for the GSEA permutations to ensure reproducibility of the analysis. Normalized enrichment scores (NES) and FDR-adjusted p-values were used to score pathways. Pathway activity inference for ‘Hypoxia’ was conducted using the Pathway RespOnsive GENes for activity inference (PROGENy) method^96^ in decoupleR. Using the top 500 most responsive genes within the Hypoxia module, pseudobulk profiles were normalized (counts per million) and analyzed using a multivariate linear model to infer pathway activity, with higher scores representing increased pathway activity. Comparisons were made among CD4 T cell subsets between CI-Sep and NHC. Pathway activity analysis for custom gene sets was performed using the AUCell method^97^ in decoupleR. Custom pathway gene sets (e.g. Sadik 2020 Pan-Tissue AHR Signature^61^) were evaluated in CD4 T cell subsets between CI-Sep and NHC. Pathway activity scores were computed based on the expression of at least 5 genes from the pathway using a random seed (28), with higher scores representing increased pathway activity.

### Flux Balance Analysis (FBA)

FBA was conducted using the Compass package^53^. For each condition (CI-Sep and NHC) and CD4 T cell subset, single-cell transcriptomic data were normalized using scanpy.pp.normalize_total as counts per million and aggregated into tab-delimited files. Compass using the IBM CPLEX optimization engine was run using the Recon2 consensus metabolic reconstruction and default settings (penalty-diffusion knn, lambda 0, num-neighbors 30) with microclusters of 25 cells. Raw reaction penalty scores were output and negative log transformed to give values representing reaction flux potential, with higher scores indicating higher reaction activity. For analysis, low-confidence reactions and reactions without variability across conditions were excluded. For bidirectional reactions, all reactions were included in the modeling but only positive fluxes were retained for visualization. Reaction activity was standardized as z-scores or calculated as Cohen’s D effect sizes for visualization and comparison between conditions.

### Statistics

All statistical analyses were performed using Python (v3.9) with scipy.stats (v1.11.4), scikit-posthocs (v0.9.0), and statsmodels (v0.14.1). Group comparisons were conducted using Mann-Whitney U tests for two-group comparisons and Kruskal-Wallis tests for multi-group comparisons, with the implementation of post-hoc pairwise comparisons via Dunn’s test. Spearman correlations were used to assess associations between continuous variables. For patient characteristics, summary statistics including medians, interquartile ranges (IQR), and p-values were calculated using the tableone package (v0.9.1).

For scRNA-seq analyses, default statistical methods were used for each package, including an empirical Bayes approach for proportionality analysis in Scanpro, the Wald test for differential gene expression in DESeq2, and a Kolmogorov-Smirnov-like statistic with permutation-based significance testing for gene set enrichment analysis (GSEA). For Compass, statistical significance was assessed using the Wilcoxon rank-sum test with Benjamini-Hochberg (B-H) adjustment for multiple comparisons. Effect sizes were represented by Cohen’s d, which was calculated using the pooled standard deviation.

Where indicated, we corrected for multiple comparisons using the B-H adjustment. Actual p-values were reported in figures unless constrained by physical space. Error bars in figures represent standard error of the mean (SEM) or 95% confidence intervals, as indicated in the figure legends.

### Data and Code Availability

Data and code are available upon request to facilitate transparency and collaboration. Full datasets, analytical code, and scRNA-seq files will be made publicly available by completion of peer review.

## ACKNOWLEDGEMENTS

We thank the patients for their participation and invaluable donations of the human tissues used in this study. We are grateful for the contributions of all clinical staff that interfaced with our research team to support this work. We appreciate the helpful discussions with Dr. Stokes Peebles, Dr. Tim Blackwell, Dr. Anna Hemnes, Dr. Henrique Serezani, Dr. Jonathan Kropski, Dr. Denis Mogilenko, and the members of the laboratories of Dr. Jeffrey Rathmell, Dr. Julie Bastarache, and Dr. Ware. We are grateful for the assistance of Dr. Angela Jones and staff in the Vanderbilt Technologies for Advanced Genomics core.

---

**Extended Data Figure 1:**
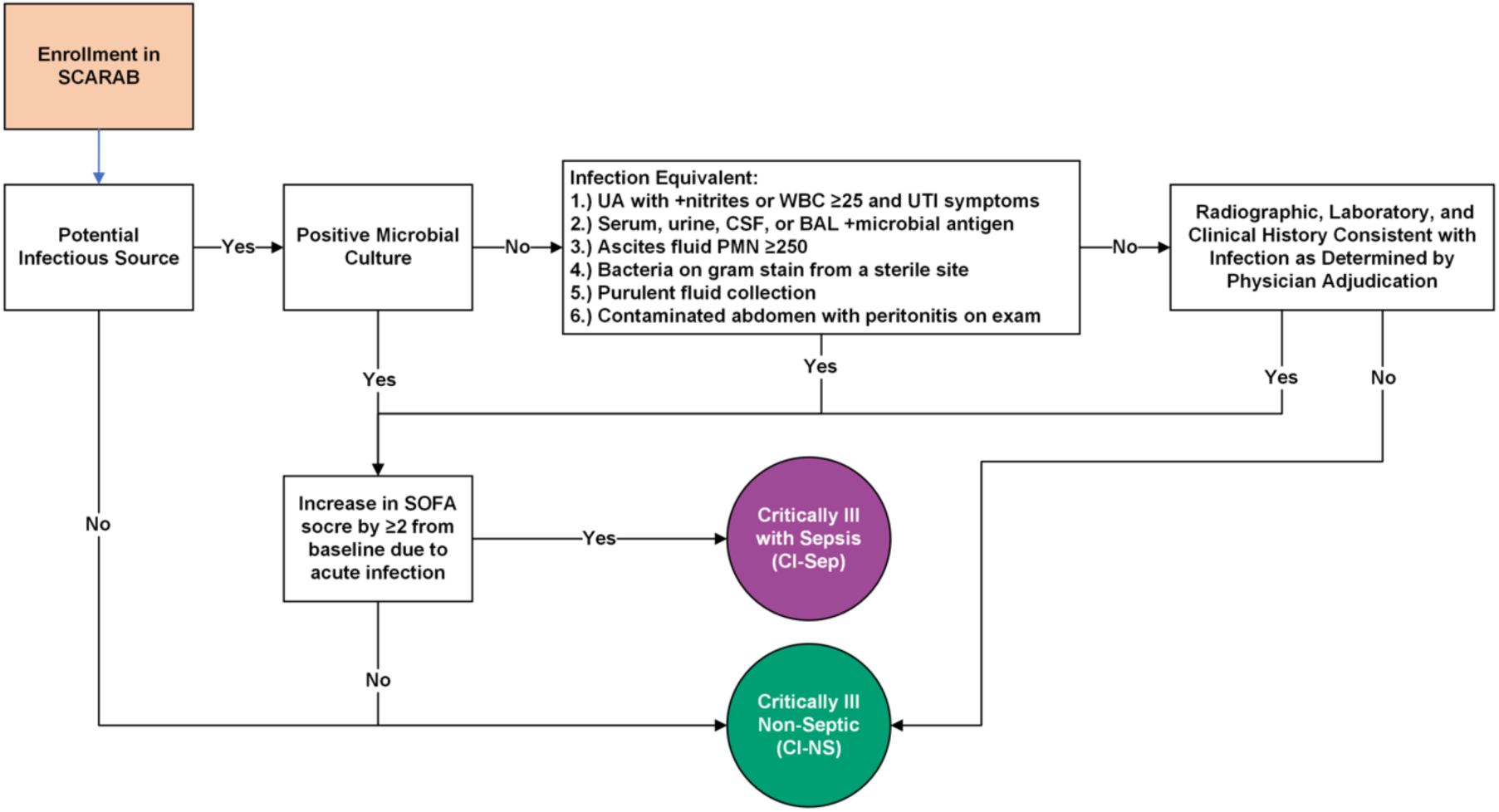
Sepsis Adjudication Algorithm.

**Extended Data Figure 2:**
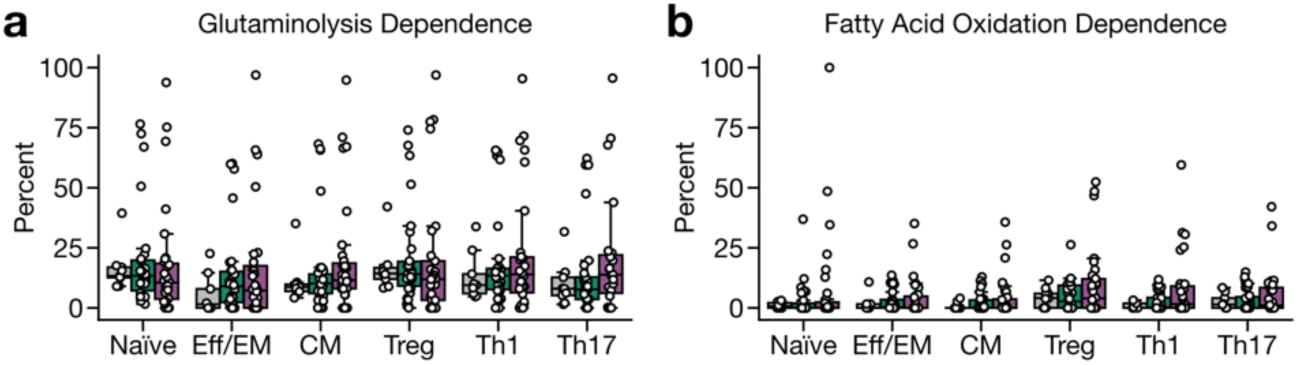
Additional Metabolic Dependencies of CD4 T Cells in Critical Illness and Severe Sepsis. **a,** Glutaminolysis and **b,** fatty acid oxidation dependence per CD4 T cell subset determined by puromycin-incorporation assay at day 2 post-ICU admission. Displayed as percentage of the maximum puromycin incorporation without metabolic inhibitors. Statistical analysis was performed using the Kruskal-Wallis test with Dunn’s post-hoc testing. Error bars represent 95% confidence intervals.

**Extended Data Figure 3:**
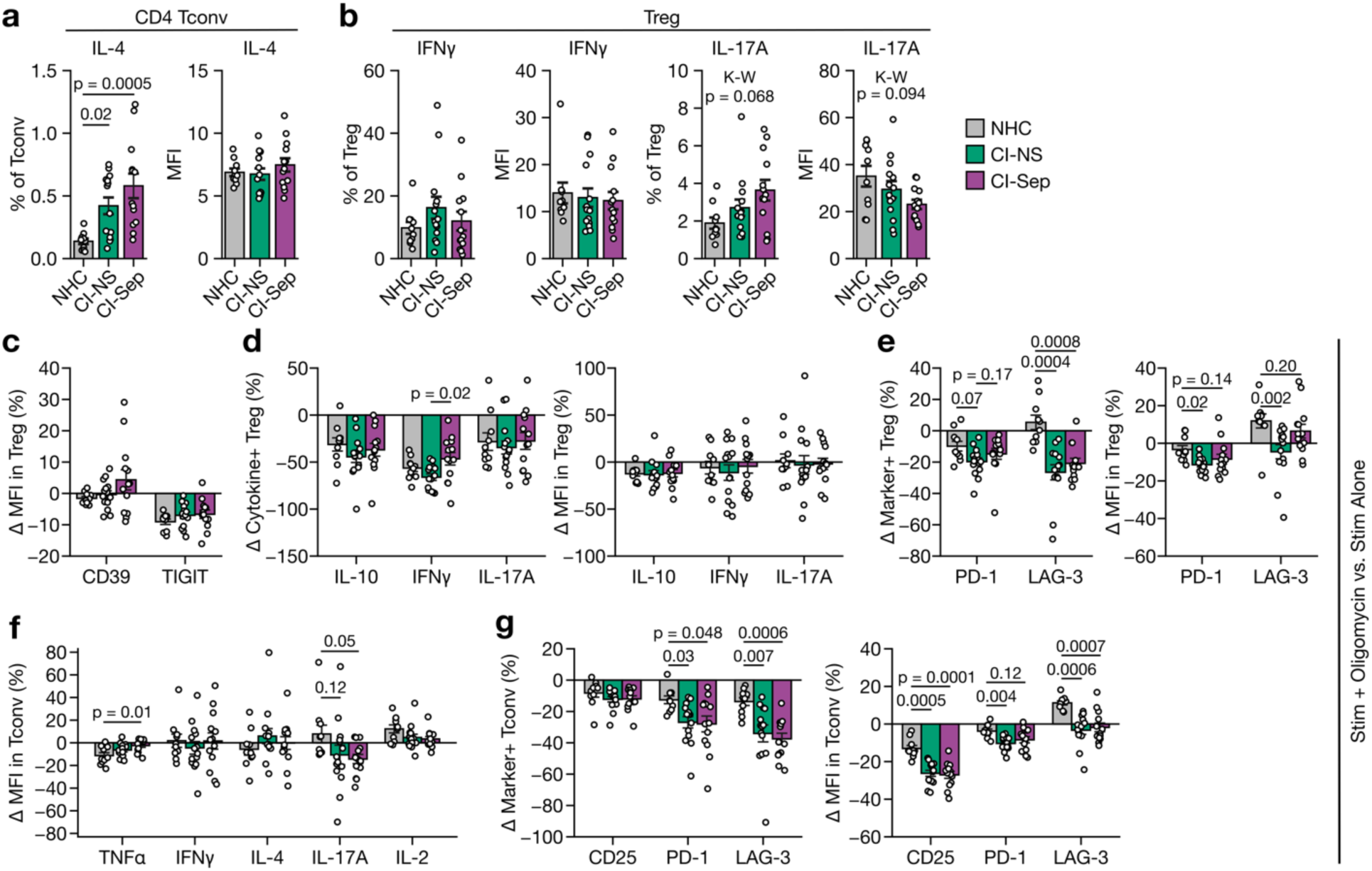
Additional functionalities of CD4 T cells experiencing mitochondrial stress. **a,** IL-4 expression in Tconv (CD4+ FOXP3-) of PBMC restimulated for 24 hrs with anti-CD3/CD28/CD2. **b,** IFNγ and IL-17A expression in Treg (CD4+ FOXP3+) of PBMC restimulated for 24 hrs with anti-CD3/CD28/CD2. **c,d,e,** Percentage change in the cytokine or marker MFI, frequency of cytokine expressing, frequency of marker expressing Treg among CD4+ T cells in anti-CD3/CD28/CD2 restimulated PBMC with stimulation alone or with stimulation plus sublethal mitochondrial disruption with oligomycin (10nM). **f,g,** Percentage change in the cytokine or marker MFI or frequency of marker expressing Tconv in anti-CD3/CD28/CD2 restimulated PBMC with stimulation alone or with stimulation plus sublethal mitochondrial disruption with oligomycin (10nM). Statistical analysis was performed using the Kruskal-Wallis test followed by Dunn’s post-hoc testing. Error bars represent SEM. K-W = Kruskal-Wallis; MFI = mean fluorescence intensity (geometric).

**Extended Data Figure 4:**
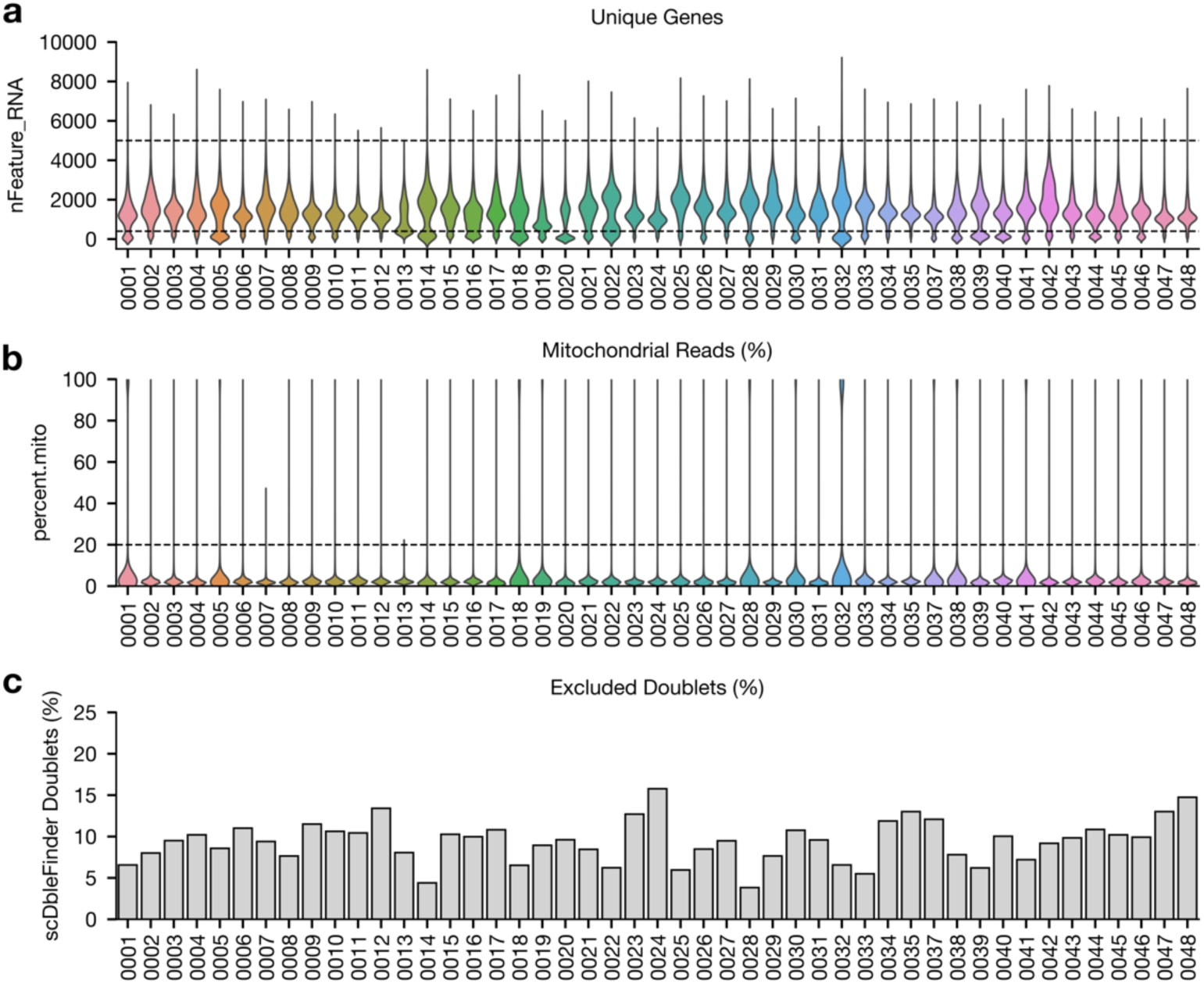
scRNA-seq quality control metrics. **a,** Number of unique genes (features) per cell after artifact and ambient RNA removal with CellBender. Dashed lines represent subsequent filtering thresholds (droplets with <400 or >5000 features were excluded). **b,** Percentage of mitochondrial reads per cell after artifact and ambient RNA removal with CellBender. Dashed line represents subsequent filtering threshold (droplets with >20% mitochondrial reads were excluded). **c,** Frequency of doublets that were removed with scDblFinder. Each sequenced sample is listed along the x-axis.

**Extended Data Figure 5:**
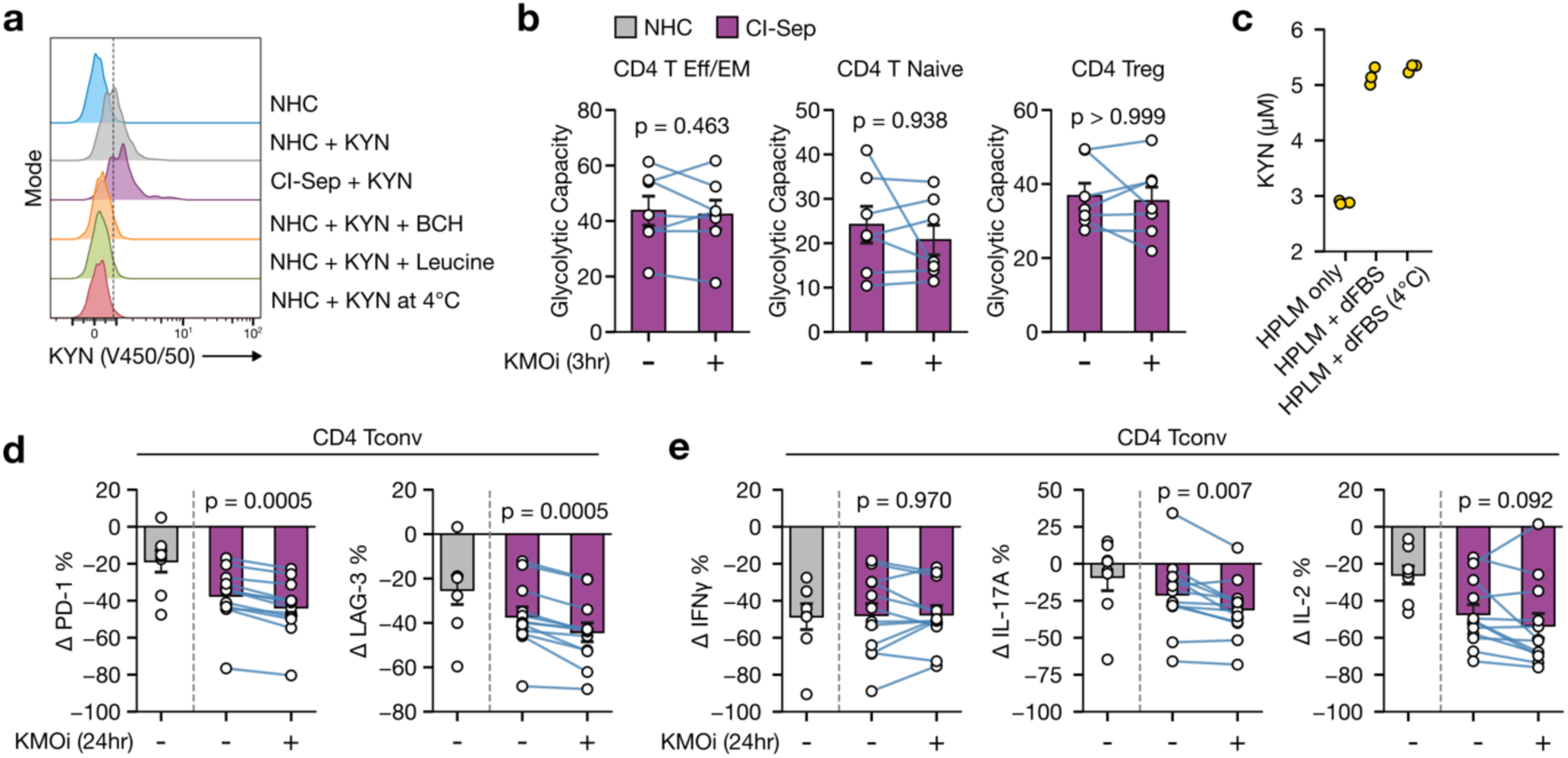
Additional effects of kynurenine metabolism modulation in CD4 T cells from critically ill septic patients. **a,** Representative flow cytometry histograms of kynurenine (KYN) uptake via fluorescence detection (450 nm filter with 50 nm bandwidth following violet laser excitation) after KYN-pulse. Controls include SLC7A5 inhibitor BCH, saturating concentration of an alternative SLC7A5 substrate leucine, and KYN pulse at 4°C. **b,** Glycolytic capacity measured by puromycin incorporation assay of paired samples. PBMC were pre-treated for 3 hrs in media alone or media plus 10 µM GSK180, a kynurenine 3-monooxygenase inhibitor (KMOi). **c,** LC-MS/MS measurement of KYN concentration in fresh HPLM, fresh HPLM with 10% dFBS, or HPLM with 10% dFBS stored at 4°C for 30 days. Percentage change in the frequency of PD-1 or LAG-3-expressing CD4 Tconv in anti-CD3/CD28/CD2 restimulated PBMC with stimulation plus oligomycin or stimulation plus oligomycin and KMOi compared to stimulated-only. **d,** Percentage change in the frequency of PD-1 or LAG-3-expressing CD4 Tconv in anti-CD3/CD28/CD2 restimulated PBMC with stimulation plus oligomycin or stimulation plus oligomycin and KMOi compared to stimulated-only. **e,** Percentage change in the frequency of cytokine-expressing CD4 Tconv in anti-CD3/CD28/CD2 restimulated PBMC with stimulation plus oligomycin or stimulation plus oligomycin and KMOi compared to stimulated-only. NHC cohort is shown in **d,e** as a reference. Each data point in **b,d,e** represents a biologic replicate, and each data point in **c** represents a technical replicate. Statistical analysis for **b,d,e** was performed using paired sample testing with Wilcoxon signed-rank test. Error bars represent SEM.

**Extended Data Figure 6:**
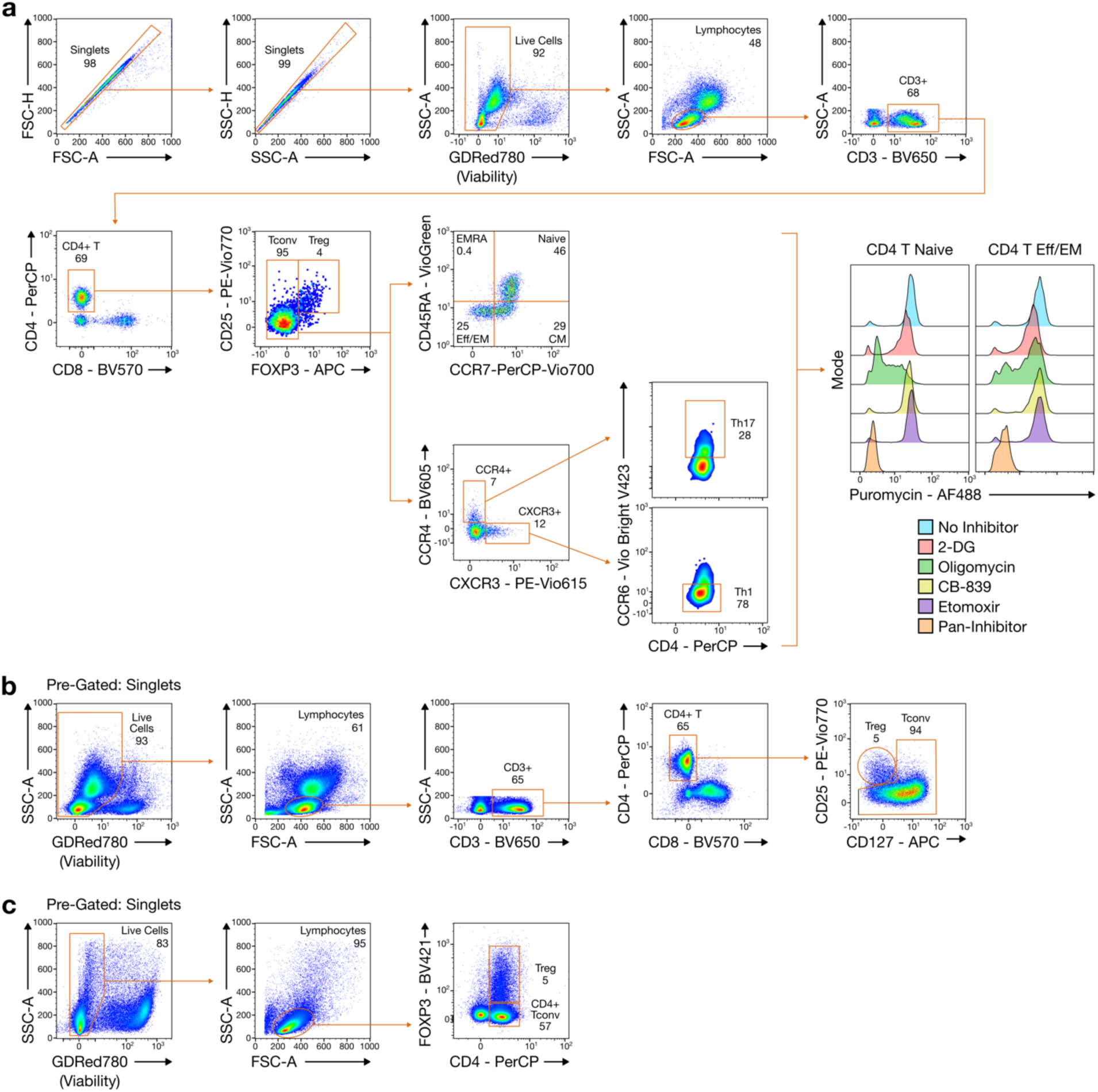
Representative flow cytometry gating strategies. **a,** Gating strategy for the identification of CD4 T naïve, Eff/EM, CM, Th1, Th17, and Treg via fixation and permeabilization. This example includes representative anti-puromycin staining. **b,** Gating strategy for identification of conventional CD4 T cells and Treg in unfixed cells using the CD25/CD127 convention. **c,** Gating strategy for the identification of CD4 Tconv and Treg in anti-CD3/CD28/CD2 restimulation experiments.

## Graphical Summary

Figure created in BioRender.com.

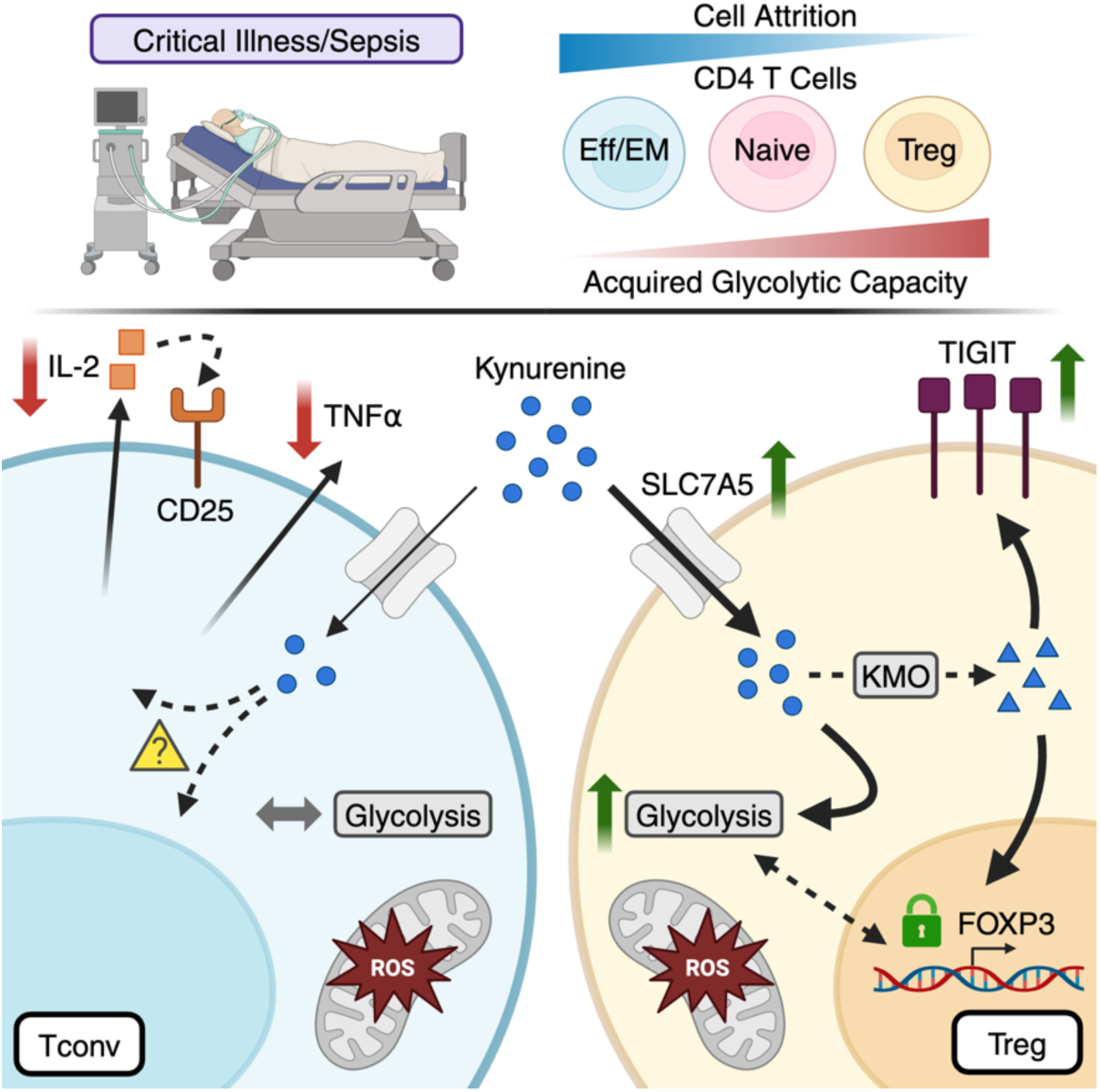

